# Crops under continuous cultivation exhibit less plasticity than those under interrupted cultivation

**DOI:** 10.1101/2024.12.10.625649

**Authors:** Ganesh Alagarasan, Rajeev K. Varshney, Eswarayya Ramireddy

## Abstract

Evolutionary studies indicate that species in stable environments often evolve with reduced plasticity, whereas those in variable environments tend to maintain higher plasticity to adapt to changing conditions. Our study explores whether this evolutionary principle extends to cultivated crops. In crop science, phenotypic plasticity is generally understood as a short-term response to environmental factors. Yet, the long-term evolutionary changes in both plastic and non-plastic traits under different cultivation regimes remain largely unexamined. Herein, we developed a novel mechanistic crop growth model, collectively termed the Trait-Environment Fitness Interaction (TEFI) Model, to study if and how trait plasticity varies among crops under different cultivation regimes. Our results, based on the TEFI Model, show higher trade-offs between fitness and plasticity. Specifically, we observed the evolution of higher plasticity in crops subjected to intermittent cultivation, which experienced more variable environments. However, this higher plasticity does not compensate for fitness losses due to the high rate of environmental unpredictability. Conversely, species under relatively stable conditions tend to evolve with reduced plasticity. Using real-world crop datasets, we validated the theoretical predictions of the TEFI Model, which suggest that the longer the interruption, the higher the plasticity. Our results highlight the evolutionary impact of cultivation patterns on trait plasticity and its importance in crop fitness. Ultimately, our findings illustrate how evolutionary principles of plasticity, as captured by the TEFI Model, can inform sustainable crop improvement strategies.

## Introduction

Phenotypic plasticity refers to the ability of living organisms to alter their phenotype in response to environmental conditions. In agricultural science, plasticity is widely documented in many important traits across different crop plants (Rahman et al., 2019; Giordano et al., 2024; Guo et al., 2024). The impact of this plasticity on agronomically desirable outcomes, however, is highly context-dependent. For example, plasticity allows roots to grow deeper in search of water, which can help crop plants avoid yield loss due to water stress (Sandhu et al., 2016; Kadam et al., 2017; Schneider et al., 2020). In contrast, plasticity exhibited by shoots during the shade avoidance response (SAR) results in taller shoots, which negatively affects crop yield (Adjesiwor et al., 2021; Qin et al., 2023; Tian et al., 2024; Gawinowski et al., 2024). Despite the potential benefits of plasticity in certain contexts, in agricultural settings, trait stability is often preferred over plasticity because the unpredictable nature of plastic traits can make it challenging to optimize crop cultivation practices. For this reason, most crop improvement programs aim to select for traits with little to no plasticity to ensure crops perform consistently in varied environmental conditions (Han et al., 2020; Ibrahim et al., 2024; Corlouer et al., 2024). As a consequence of this focus on stability, relatively little attention has been given to understanding how trait plasticity evolves under agricultural conditions. With an unpredictably changing environmental conditions and improving farming methods, studying phenotypic plasticity has become more important for making crops resilient and increasing agricultural productivity (Kusmec et al., 2018; Schneider and Lynch, 2020; Monforte, 2020; Laitinen, 2024). Although trait stability has traditionally been prioritized in breeding programs, recent research has uncovered a distinct genetic architecture governing both trait ‘mean’ and trait ‘plasticity’ (Tetard-Jones et al., 2011; Kusmec et al., 2017; Li et al., 2018; Jin et al., 2023; Chen et al., 2024). These findings are important because they suggests that plasticity and the average expression of traits are controlled by different genetic factors, opening up new possibilities for selective breeding. This genetic insights enables breeders to more precisely target and manipulate either trait stability or plasticity, depending on the environmental conditions or agricultural goals. As a result, it could pave the way for developing crops that are not only high-performing under ideal conditions but also resilient to fluctuating environments. Despite these advancements in understanding the genetic basis of plasticity, significant gaps remain, particularly regarding how plasticity evolves under agricultural conditions over time. To address this gap, we have developed a novel Trait-Environment Fitness Interaction (TEFI) Model to study how plasticity evolves under different cultivation practices, including continuous cultivation (similar to modern crops) and interrupted cultivation (resembling underutilized crops). Contrary to the traditional crop growth models, which predict crop suitability based on environmental factors (Pironon et al., 2019; Zonneveld et al., 2023), the TEFI model mechanistically explores how trait evolves under environmental stochasticity. Alongside this, the TEFI model also predicts long-term crop fitness. Our findings, supported by real-world agricultural datasets, suggest that crops under continuous cultivation evolve less plasticity compared to those under interrupted cultivation. More broadly, we observe that environmental stochasticity can reduce the benefits of plasticity and increase the risk of extinction, underscoring the importance of incorporating these dynamics into crop breeding strategies.

## Materials and methods

Let us consider five different types of crop species based on their cultivation patterns: continuous cultivation, short-interrupted cultivation, mid-interrupted cultivation, long-interrupted cultivation, and noncontinuous interrupted cultivation. These patterns reflect various strategies that crops may follow due to environmental and human factors (due to focus on primary food crops). In particular, the interruption periods are designed to mimic the cultivation of underutilized crops, which often experience gaps in cultivation. Using the TEFI model, we model how fitness is affected by a stochastic environment to understand the long-term viability of these crops and how they may respond to unpredictable variations in conditions such as temperature. Below, we will explain the specific equations within the TEFI model used for fitness estimation and the rules governing each scenario.

### (a) Fitness dynamics under different cultivation and environmental variation

#### (i) Fitness dynamics under continuous cultivation and environmental variation

In continuous cultivation, species are cultivated without any interruptions. This pattern is typical of modern staple crops like rice, wheat, and maize, planted yearly without fallow periods. These species are continuously exposed to environmental conditions, adapting over time to predictable variations such as temperature. For these species , the fitness at year *t* can be determined using the TEFI base fitness equation:

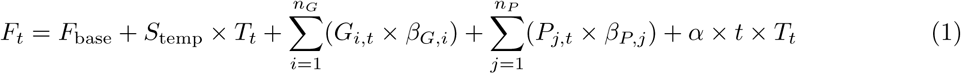

The fitness of a species in any given year, denoted as *F_t_*, is influenced by multiple factors, which are captured in the equation provided. Here, *F*_base_ represents the base fitness of the species, essentially its viability without any external influences. This foundational fitness level is then modified by a series of terms that account for environmental and trait-based effects. The temperature sensitivity coefficient, *S*_temp_, is multiplied by the temperature at year *t* (*T_t_*), representing how changes in temperature directly influence fitness. This term reflects the species’ immediate response to environmental conditions, acknowledging that temperature variations can either enhance or diminish fitness depending on the species’ sensitivity. Non-plastic traits, which are inherited characteristics that do not change in response to the environment, contribute to fitness through the term 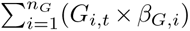. Each non-plastic trait *G_i,t_* affects fitness based on its specific effect size *β_G,i_*, and the sum across all non-plastic traits (*n_G_*) provides a cumulative impact on fitness. These traits represent stable genetic contributions that shape fitness regardless of external conditions. Conversely, plastic traits, which can change in response to environmental factors, contribute through the term 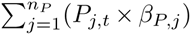. Each plastic trait *P_j,t_* influences fitness depending on its own effect size *β_P,j_*, allowing for phenotypic adjustments that help the species cope with environmental variations. The effect sizes for both genetic (*β_G,i_*) and plastic (*β_P,j_*) traits are modeled as random variables drawn from normal distributions, specifically *β_G,i_* ∼ *N* (0, 0.1) for genetic traits and *β_P,j_*∼ *N* (0, 0.1) for plastic traits, where the mean is 0 and the variance is 0.1. The total contribution from these traits, summed across *n_P_*plastic traits, reflects the species’ ability to adapt phenotypically to its surroundings. In simulation, the update rules for each trait depend on whether the trait is plastic or non-plastic. For non-plastic traits, the trait value at the next time step (*t* + 1) is updated by adding the change due to mutation (*δ*_mutation_) to the current trait value (*t*), represented as Trait*_t_*_+1_ = Trait*_t_* + *δ*_mutation_. In contrast, for plastic traits, the update incorporates both mutation and plasticity effects, calculated as Trait*_t_*_+1_ = Trait*_t_* + *δ*_mutation_ + *δ*_plasticity_. The mutation change (*δ*_mutation_) occurs with a probability *µ* = 0.005, and when mutation occurs, the change is drawn from a normal distribution with mean 0 and variance 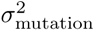, where *σ*_mutation_ = 0.05. If mutation does not occur, *δ*_mutation_ = 0. For plastic traits, the plasticity change (*δ*_plasticity_) occurs with a probability *ρ* = 0.01, and when it occurs, the change is drawn from a normal distribution with variance 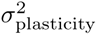, where *σ*_plasticity_ = 0.1. If plasticity does not occur, *δ*_plasticity_ = 0. Both mutation and plasticity effects are constrained to ensure that trait values remain within the range of 0 to 1. The adaptation term *α*×*t*×*T_t_* encapsulates the long-term evolutionary adaptation of the species. This term links the passage of time (*t*) and environmental temperature (*T_t_*) to gradual improvements in fitness, representing how species evolve over time to become more suited to their environments. The rate of this adaptation is controlled by the coefficient *α*, which quantifies the speed at which fitness improves over successive years (similar to genetic gain from breeding program).

#### (ii) Fitness dynamics under different interrupted cultivation and environmental variation

The TEFI Model also accounts for interrupted cultivation patterns, allowing us to assess fitness dynamics across different types of interruptions. The interruptions in cultivation are classified into four distinct types. A short interruption occurs over 25% of the total years of simulation. A mid-length interruption lasts for 50% of the total years, while a long interruption spans 75% of the total years. A non-continuous interruption consists of randomly generated interruption periods. For species subjected to interrupted cultivation, fitness is calculated across three distinct phases: before, during, and after the interruption. Prior to the interruption, fitness is determined following the approach outlined in Equation 1. Fitness during the interruption period (when *t*_start_ ≤ *t* ≤ *t*_end_) is calculated using the following equation:

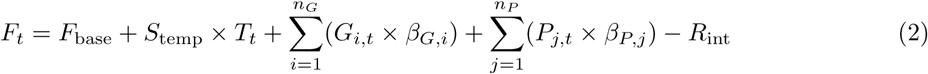

Here, *R*_int_ is the reduction factor during the interruption, which quantifies the decrease in fitness due to the interruption. It is calculated based on the temperature difference between the start and end of the interruption period, scaled by a constant *k*:

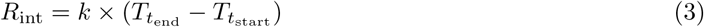

Here, *k* is a constant representing the proportion of the temperature difference contributing to fitness reduction (e.g., *k* = 0.3 or *k* = 0.5), *T_t_*_start_ and *T_t_*_end_ are the temperatures at the start and end years of the interruption, respectively. During periods of cultivation interruption, species face adverse conditions that can negatively impact their fitness. The fitness equation for this period modifies the continuous cultivation equation by introducing the reduction factor *R*_int_, which directly subtracts from the fitness. This term represents the immediate negative impact of the interruption on the species’ fitness. By calculating *R*_int_ based on temperature differences, the model captures the idea that significant environmental changes during the interruption exacerbate its negative effects. A larger temperature difference implies greater environmental instability, which can further reduce fitness.

After the interruption period (when *t > t*_end_), the fitness is calculated by adjusting the base fitness and trait contributions to account for the cumulative impact of the interruption. The fitness at year *t* is determined using the following equation:

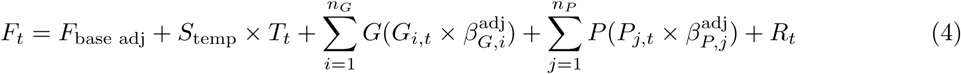

The fitness recovery model incorporates the impact of cultivation interruptions by adjusting both the base fitness and the contributions of genetic and plastic traits over time. When an interruption occurs, a cumulative interruption impact *C*_impact_ accumulates, representing the negative effects of the disruption. This impact grows incrementally as the interruption continues, modeled by the equation *C*_impact_ = *C*_impact_ + *R*_int_ × *D*_int_, where *R*_int_ is the reduction factor during the interruption, and *D*_int_ is the duration of the interruption in years. As the cumulative impact increases, both the base fitness *F*_base_ and the effect sizes of the non-plastic and plastic traits (*β_G,i_* and *β_P,j_*, respectively) are scaled down, capturing the lasting damage caused by the interruption. These adjustments are expressed in the following equations: 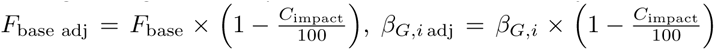, and 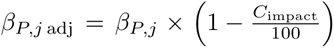. These adjustments represent how the species’ baseline fitness and the effectiveness of its traits decrease as the interruption progresses. Following the end of the interruption, species begin the process of recovery, gradually improving their fitness. The recovery is modeled by the term *R_t_* = *r*_adj_ × (1 − *F_t_*), where *R_t_* represents the amount of fitness regained at year *t*, depending on the current fitness deficit (1 − *F_t_*) and the adjusted recovery rate *r*_adj_. The adjusted recovery rate is influenced by the duration of the interruption, expressed as 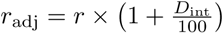, where *r* is the base recovery rate. Longer interruptions slow down the recovery process, as reflected by this equation, because species face greater challenges in rebuilding their fitness after prolonged disruptions.

### (b) Quantitative assessment of trait evolution dynamics

The evolution of plastic and non-plastic traits is estimated over time using a matrix of average trait values, *T* ∈ R*^n^*^years^ *^×n^*^traits^ , where *T_i,j_* represents the average value of trait *j* in year *i*. The average value *T_i,j_* is calculated as 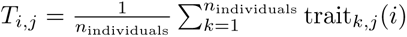. This approach enables the tracking and quantification of how traits evolve in response to different environmental conditions, such as continuous cultivation, and interruptions of varying durations (short, mid, long, and non-continuous).

### (c) Model testing via simulation

The simulations were conducted in Python (version 3.11.4) using the parameters listed in Table 1. The TEFI Model’s complexity, including the dynamic interactions between genetic traits, plastic traits, and various environmental interruptions, was carefully coded to ensure accurate behavior and performance (refer File S1 and S2 for Pyhton codes). Using NumPy for numerical computations and Pandas for data management, the simulation tracked changes in traits and fitness over time. The TEFI Model simulated multiple interruption scenarios, including continuous cultivation and intermittent cultivation (short, mid, long, and non-continuous), with interruptions affecting fitness and recovery over time. Fitness was computed annually based on the influence of both non-plastic and plastic traits, as well as environmental factors like temperature. In our study, we utilized temperature anomalies for the period 1851 to 2022 (NOAA, 2023), referenced against the 1901-2000 average, to predict species responses (Fig. S1A, Additional Sheet 1). Vinton et al. (2022) emphasized the significant impact of environmental condition variations and their predictability on the evolutionary pathways available to organisms, particularly through phenotypic plasticity and genetic adaptation. In our study, we extend these considerations by examining the rate of change, variance, and temporal autocorrelation in temperature anomalies. If the temperature and its nature of change were to remain constant, we might not expect to see significant shifts in the model outcomes. The upward trend in temperature anomalies over time (Fig. S1A), along with the rate of change (Fig. S1B), the variance in anomalies (Fig. S1C), and the autocorrelation at various lags (Fig. S1D), render this dataset robust for predicting evolutionary responses to climate change and cultivation practices.

**Table 1.** List of parameters used in the TEFI model simulation.

| Parameter | Description | Value |
| --- | --- | --- |
| $F_{\text{base}}$ | Base fitness of the species without external influences. | 0.5 |
| $S_{\text{temp}}$ | Temperature sensitivity coefficient. | -0.02 |
| $T_t$ | Temperature at year $t$. | From temperature data |
| $G_{i,t}$ | Non-plastic trait $i$ at year $t$. | Random values between 0 and 1 |
| $\beta_{G,i}$ | Effect size of non-plastic trait $G_{i,t}$, drawn from $\mathcal{N}(0, 0.1)$. | From $\mathcal{N}(0, 0.1)$ |
| $P_{j,t}$ | Plastic trait $j$ at year $t$. | Random values between 0 and 1 |
| $\beta_{P,j}$ | Effect size of plastic trait $P_{j,t}$, drawn from $\mathcal{N}(0, 0.1)$. | From $\mathcal{N}(0, 0.1)$ |
| $\alpha$ | Adaptation rate. | 0.001 |
| $\mu$ | Probability of mutation for both plastic and non-plastic traits per year. | 0.005 |
| $\rho$ | Probability of plasticity change for plastic traits per year. | 0.01 |
| $\sigma_{\text{mutation}}$ | Standard deviation of the mutation effect on traits. | 0.05 |
| $\sigma_{\text{plasticity}}$ | Standard deviation of plasticity effect on plastic traits. | 0.1 |
| $\delta_{\text{mutation}}$ | Change due to mutation. | Random, based on mutation probability |
| $\delta_{\text{plasticity}}$ | Change due to plasticity. | Random, based on plasticity probability |
| $t$ | Year (time step) in the simulation. | 0 to $n_{\text{years}}$, $n = 172$ |
| $n_G$ | Number of non-plastic traits. | 5 |
| $n_P$ | Number of plastic traits. | 5 |

### (d) Parameter sensitivity analysis

To evaluate the robustness of the TEFI Model and the impact of key parameters on fitness under different scenarios, we conducted a global sensitivity analysis (GSA) using Saltelli sampling and Sobol analysis (refer File S3 for Python codes). The sensitivity analysis aimed to quantify how variation in mutation rates, plasticity changes, temperature sensitivity, and adaptation rates impacted average fitness across several interruption scenarios, including continuous cultivation, short, mid, long, and non-continuous interruptions. Seven key parameters were varied within predefined ranges: mutation rate (0.001 - 0.01), mutation effect (0.01 - 0.1), plasticity rate (0.001 - 0.05), plasticity effect (0.05 - 0.2), temperature sensitivity (-0.05 - 0.01), adaptation rate (0.0001 - 0.005), and recovery rate (0.001, 0.02). Saltelli’s method was used to sample the parameter space, generating 1000 samples per parameter, resulting in a total of 7000 parameter combinations, which were then used in the fitness model simulations. For each sampled parameter set, fitness was simulated across five distinct cultivation scenarios. Fitness for each scenario was calculated annually based on mutation, plasticity changes, and interruption impacts, and the average fitness over the simulation period was recorded. Sobol analysis was applied to decompose the total variance in fitness into contributions from individual parameters and their interactions. Both first-order sensitivity indices (S1), representing the direct effect of a parameter on fitness, and total-order sensitivity indices (ST), capturing direct and interaction effects, were calculated for each scenario.

### (e) Case studies in real-world agricultural datasets

#### (i) Case study in modern cereal crops - rice and wheat

We aim to investigate species-level differences in phenotypic plasticity to determine if our theoretical insights are consistent with real-world agricultural settings. For this study, we have selected two widely cultivated modern cereal crops: rice (Yano et al., 2016) and wheat (Khan et al., 2022). The rice dataset consists of 176 varieties developed through a Japanese breeding program, with phenotypic data collected for seven traits: days to heading, plant height (cm), panicle length (cm), panicle number per plant, spikelet number per panicle, leaf blade width (cm), and awn length (cm). For our analysis, we excluded the awn length trait, as it is a kind of binary trait (presence or absence), and focused on assessing plasticity in the remaining six traits. The phenotypic data were originally measured in 2013 and 2014 (representing two environments: year × location combination) for a genome-wide association (GWAS) study. For more details on crop cultivation practices and phenotypic measurements, refer to Yano et al. (2016). On the other hand, the wheat dataset consists of 280 genotypes, including a mix of advanced breeding lines and some commercial Indian varieties, with phenotypic data collected for six traits: grain filling duration (GFD), grain number per spike (GNPS), grain weight per spike (GWS), grain yield (GY), days to heading (DH), and plant height (PH). The phenotypic data were originally measured across five different locations in 2020-21 for a GWAS study. For more information on crop cultivation practices and phenotypic measurements, refer to Khan et al. (2022).

#### (ii) Case study in pulse crops - chickpea and pigeonpea

The chickpea dataset consists of 2,972 genotypes, primarily landraces, along with breeding lines and wild accessions. The phenotypic data were originally collected between 2014 and 2016, covering 12 environments (a combination of 6 locations and 2 years) for a GWAS study. Key traits measured include pods per plant (PPP), which indicates the reproductive efficiency of each genotype, hundred seed weight (HSW) as an indicator of seed size and yield potential, plant height (PH) reflecting the vegetative growth habit, days to flowering (DTF) to assess flowering time, and days to maturity (DtM) to understand the growth duration. For a full list of traits and further details on crop cultivation practices and phenotypic measurements, refer to Varshney et al. (2021). The pigeonpea dataset consists of 286 genotypes, containing landraces and breeding lines. The phenotypic data were originally collected between 2013 and 2014, covering 2 environments (a combination of 1 locations and 2 years) for a GWAS study. Key traits measured include number of pods (NP), hundred seed weight (HSW), days to flowering (DTF), plant height (PH), and days to maturity (DtM). For a full list of traits and further details on crop cultivation practices and phenotypic measurements, refer to Varshney et al. (2017).

#### (iii) Case study in underutilized crops - proso millet and foxtail millet

The proso millet dataset consists of 516 genotypes, primarily landraces, along with some breeding lines and wild accessions. The phenotypic data were originally collected between 2019 and 2020, covering 14 environments (a combination of 2 years and 7 locations), for a GWAS study. Key traits measured include panicle weight per plant, seed weight per plant, effective tillering rate, main stem length, and 1000 seed weight, among others (Additional Sheet 2). For a full list of traits and details on crop cultivation practices and phenotypic measurements, refer to Chen et al. (2023). The foxtail millet dataset consists of 680 genotypes, including a mix of landraces, wild types, and cultivated varieties. The phenotypic data were originally collected between 2010 and 2020, covering 11 environments (a combination of years and locations) for a GWAS study. Key traits measured include panicle weight per plant (PWP), grain weight per plant (GPP), plant height (PH), heading date (HD), and 1000 grain weight (1000 SW), among others. For a full list of traits and further details on crop cultivation practices and phenotypic measurements, refer to He et al. (2023).

Quantifying plasticity across diverse biological settings presents several common challenges inherent to the methods used (Valladares et al., 2006). Many approaches assume normal distribution of data, which may not align with the real-world distribution of biological traits, potentially skewing results. Additionally, the need for data to conform to specific statistical properties, such as normality or linearity, can limit the applicability of some methods to complex ecological or evolutionary scenarios where these assumptions are violated. Furthermore, the requirement for equal or substantial sample sizes across conditions in many methodologies can be a significant constraint in studies where obtaining balanced data is impractical. Here, we provide a metric to quantify plasticity offering advantages such as handling unequal sample sizes across conditions, robustness to outliers, versatility across diverse data distributions, and applicability to multivariate data, making it particularly relevant and useful in agricultural settings. It is particularly useful for meta-analyses and comparative studies that involve data from varied sources and conditions. The Plasticity Score is calculated for all the traits based on a matrix of trait values across different environmental conditions, *T* ∈ R*^nconditions×ntraits^*, where *T_i,j_* represents the value of trait *j* under environmental condition *i*. First, the median value for each trait across all environmental conditions is computed as 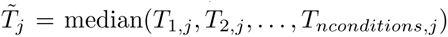, representing the central tendency of trait *j* across the environmental conditions. Next, the absolute difference between each trait value and its median is calculated as 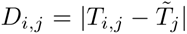. These absolute differences are then averaged over all environmental conditions for each trait, 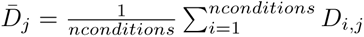, which quantifies the plasticity of trait *j* across different environmental conditions. Here, the traits with higher *D̄_j_* will have higher plasticity.

## Results

### (a) Fitness dyanmics under different cultivation scenarios

The results of the simulation indicate distinct differences in the evolutionary fitness trajectories of populations under continuous cultivation and intermittent cultivation with varying interruption types. For continuous cultivation, where fitness *F_t_* was modeled according to the TEFI Model’s base fitness equation (Eq. 1), the fitness remained relatively stable (mean = 0.497; SD = 0.149), with minimal changes over time, as indicated by the very small slope of 0.000203. This reflects slow but steady adaptation without significant disruptions (Fig. 1A). In contrast, intermittent cultivation scenarios produced more complex fitness dynamics, modeled with the Eq.2, during the interruption periods. The non-continuous interruption scenario provided a more favorable outcome for populations, as fitness before the interruption was moderate (mean = 0.451; SD = 0.206), increased slightly during the interruptions (mean = 0.522; SD = 0.167), and maintained relatively high levels after the interruptions (mean = 0.625; SD = 0.184). This suggests that the intermittent nature of the interruptions allowed organisms to periodically recover, as indicated by the adjusted recovery rate *r*_adj_, allowing them to better adapt and recover from environmental disturbances (Fig. 1B). For populations subjected to a short interruption, fitness before the interruption was moderate (mean = 0.452; SD = 0.108), but increased during the interruption (mean = 0.550; SD = 0.156), likely due to the resilience provided by plasticity traits *P_j,t_*. Post-interruption, fitness recovered strongly (mean = 0.886; SD = 0.189), as organisms were able to regain fitness quickly after a brief interruption (Fig. 2A, 2B, 2C). In the case of mid interruptions, fitness before the interruption was relatively high (mean = 0.492; SD = 0.150), but dropped during the interruption (mean = 0.415; SD = 0.146), with fitness collapsing almost completely afterward (mean = 0.0002; SD = 0.0015) (Fig. 2A, 2B, 2C). This suggests that the population’s ability to recover was severely impaired by the extended interruption, with cumulative interruption impact *C*_impact_ reducing both the base fitness *F*_base_ and the contributions of traits *β_G,i_* and *β_P,j_*. For long interruptions, fitness before the interruption was moderate (mean = 0.507; SD = 0.200), but declined sharply during the interruption (mean = 0.083; SD = 0.087), with minimal recovery afterward (mean = 0.012; SD = 0.033), indicating that the population faced extreme difficulty in rebuilding fitness after the prolonged disruption (Fig. 2A, 2B, 2C). These findings highlight that while short interruptions allowed populations to recover quickly, longer and continuous interruptions caused severe reductions in fitness, and recovery was slower or nonexistent depending on the duration and nature of the interruption. This underscores the importance of both the duration (*D*_int_) and frequency of interruptions in shaping evolutionary outcomes, with longer interruptions leading to cumulative fitness loss through *C*_impact_.

**Figure 1:**
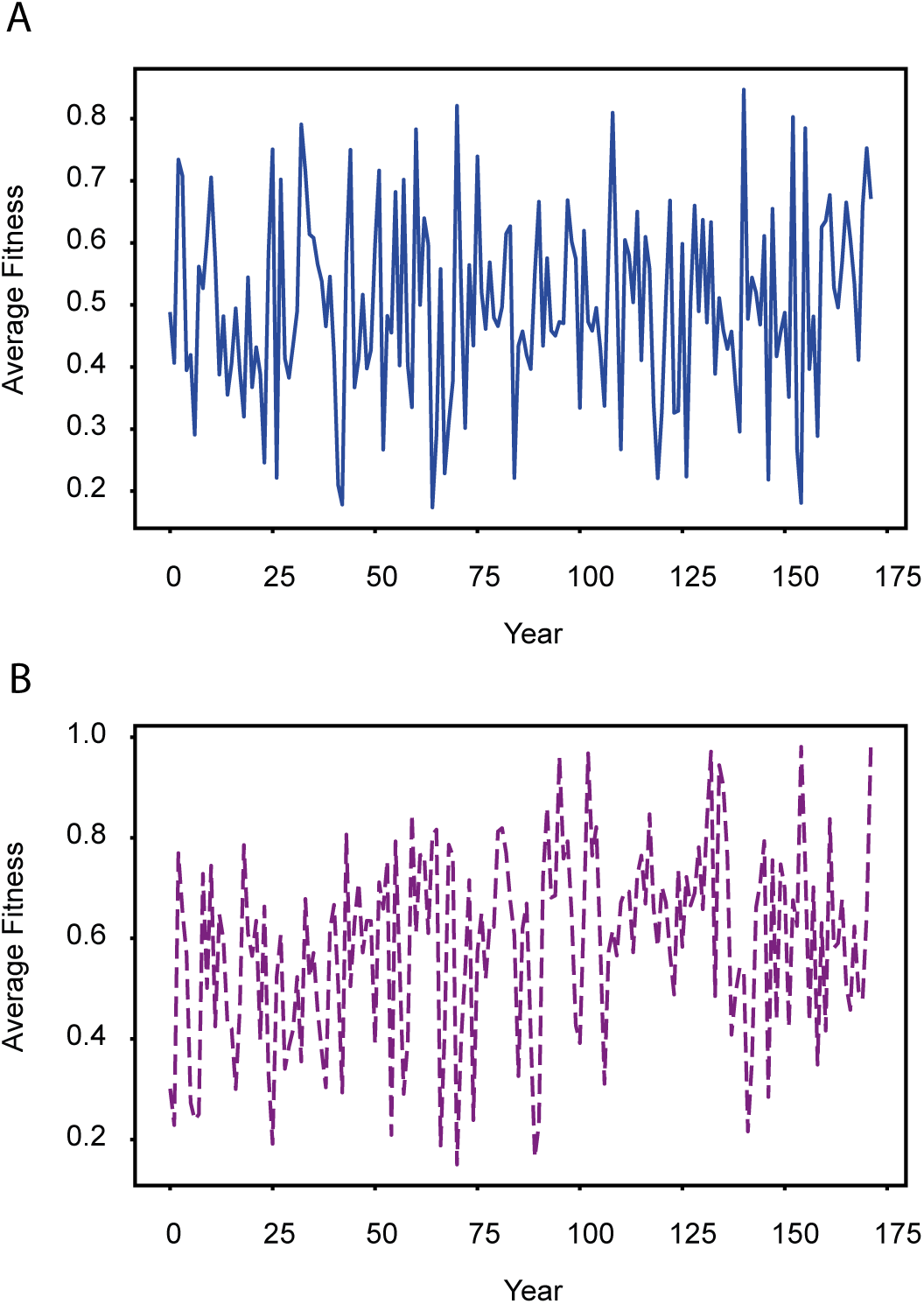
Fitness trajectories under continuous and non-continuous cultivation. (A) The fitness trajectory of species under continuous cultivation, illustrating how fitness changes over time in the absence of environmental interruptions. (B) The fitness trajectory of species under non-continuous cultivation, showing the effects of three randomly selected (refer Methods) non-continuous interruption periods on fitness, as well as the recovery patterns following these interruptions.

**Figure 2:**
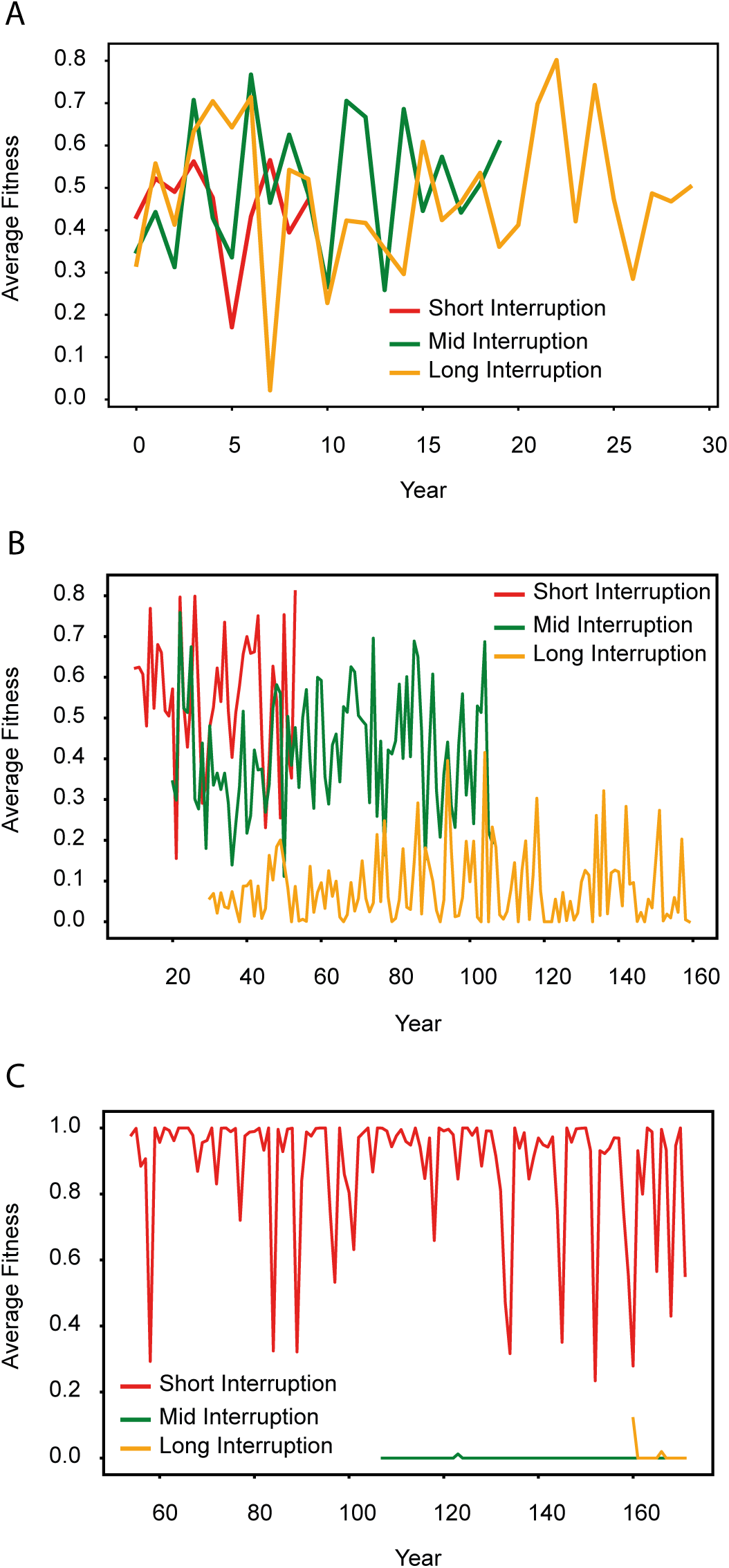
Fitness trends before, during, and after interruptions of varying durations. (A) Fitness trends before the onset of short, mid, and long interruptions. The short interruption begins in year 10, the mid interruption in year 20, and the long interruption in year 30. (B) Fitness trajectories during the interruption periods. (C) Fitness recovery patterns after the end of each interruption, highlighting differences in recovery rates following interruptions of varying durations.

### (b) Trait evolution dynamics under different cultivation scenarios

The rates of change for both genetic and plastic traits were analyzed across different environmental conditions using the TEFI Model: continuous cultivation, short interruption, mid interruption, long interruption, and non-continuous interruption. Under continuous cultivation, genetic traits exhibited a minimal positive rate of change (9.39 × 10*^−^*^6^), while plastic traits showed a slight negative trend (−7.06 × 10*^−^*^5^) (Fig. 3A). However, non-continuous interruptions led to a slight increase in genetic traits (1.52 × 10*^−^*^5^) but a decline in plastic traits (−6.71 × 10*^−^*^5^), suggesting that irregular disruptions may reduce adaptive flexibility (Fig. 3B). In scenarios with short interruptions, genetic traits displayed a minor decline (−1.51 × 10*^−^*^5^), whereas plastic traits increased (5.86 × 10*^−^*^5^), indicating an adaptive response (Fig. 4A). During mid-length interruptions, genetic traits also decreased slightly (−8.92×10*^−^*^6^), with plastic traits showing a smaller positive trend (1.36 × 10*^−^*^5^) (Fig. 4B). Long interruptions caused the most significant increases, with genetic traits increasing (2.88 × 10*^−^*^5^) and plastic traits showing the highest rate of growth (1.05 × 10*^−^*^4^), likely due to prolonged environmental stress (Fig. 4C). Overall, the results indicate that interruptions, particularly longer ones, promote plasticity, while continuous and non-continuous cultivation lead to smaller or negative changes in plastic traits.

**Figure 3:**
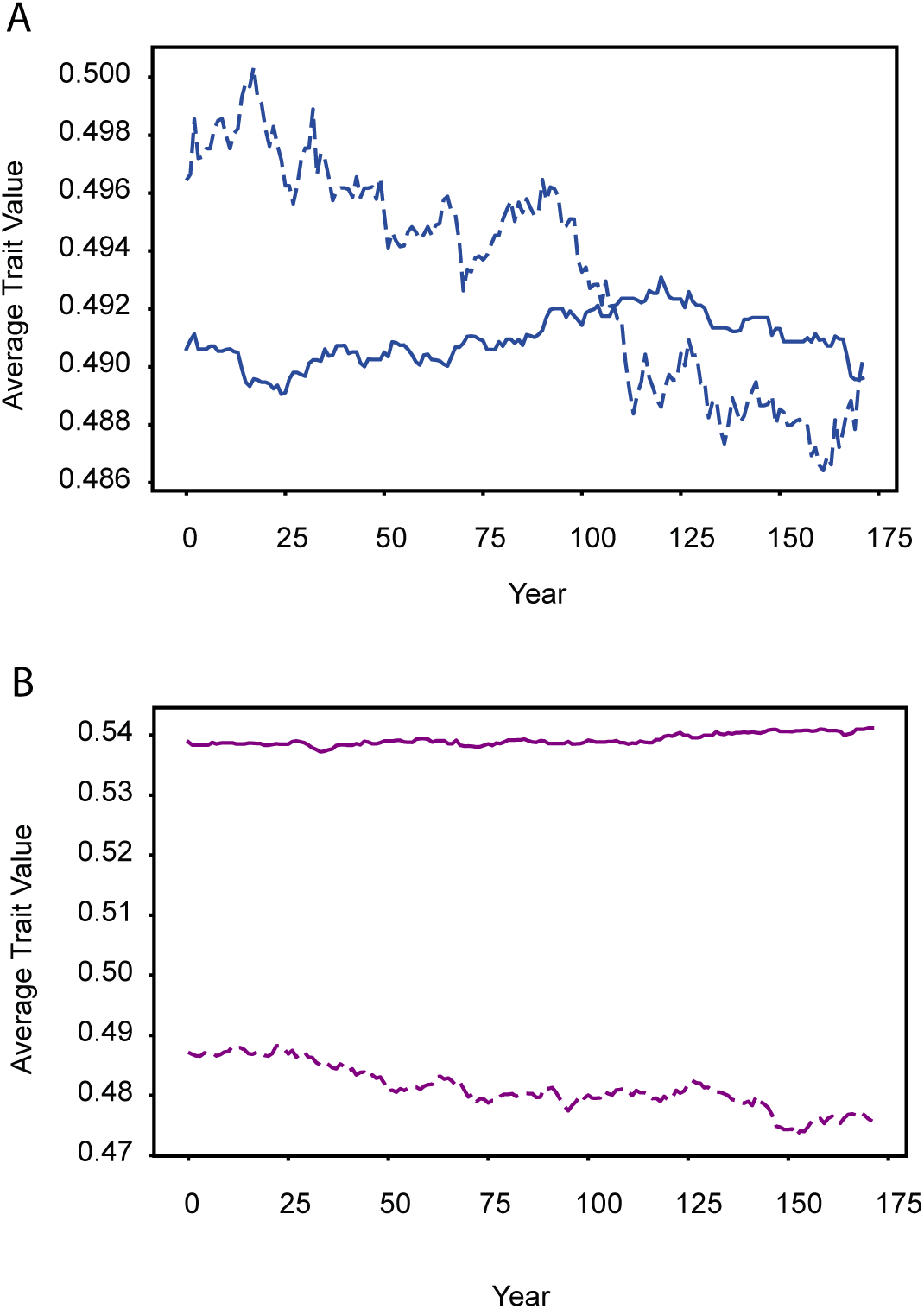
Evolution of genetic and plasticity traits under continuous and non-continuous cultivation. (A) The evolution of genetic and plasticity traits over time for species under continuous cultivation, showing how both traits change in the absence of interruptions. The solid line represents the genetic traits, while the dashed line represents the plasticity traits, illustrating their different rates of change over time. (B) The evolution of genetic and plasticity traits for species under non-continuous cultivation, depicting the effects of three randomly occurring non-continuous interruption periods.

**Figure 4:**
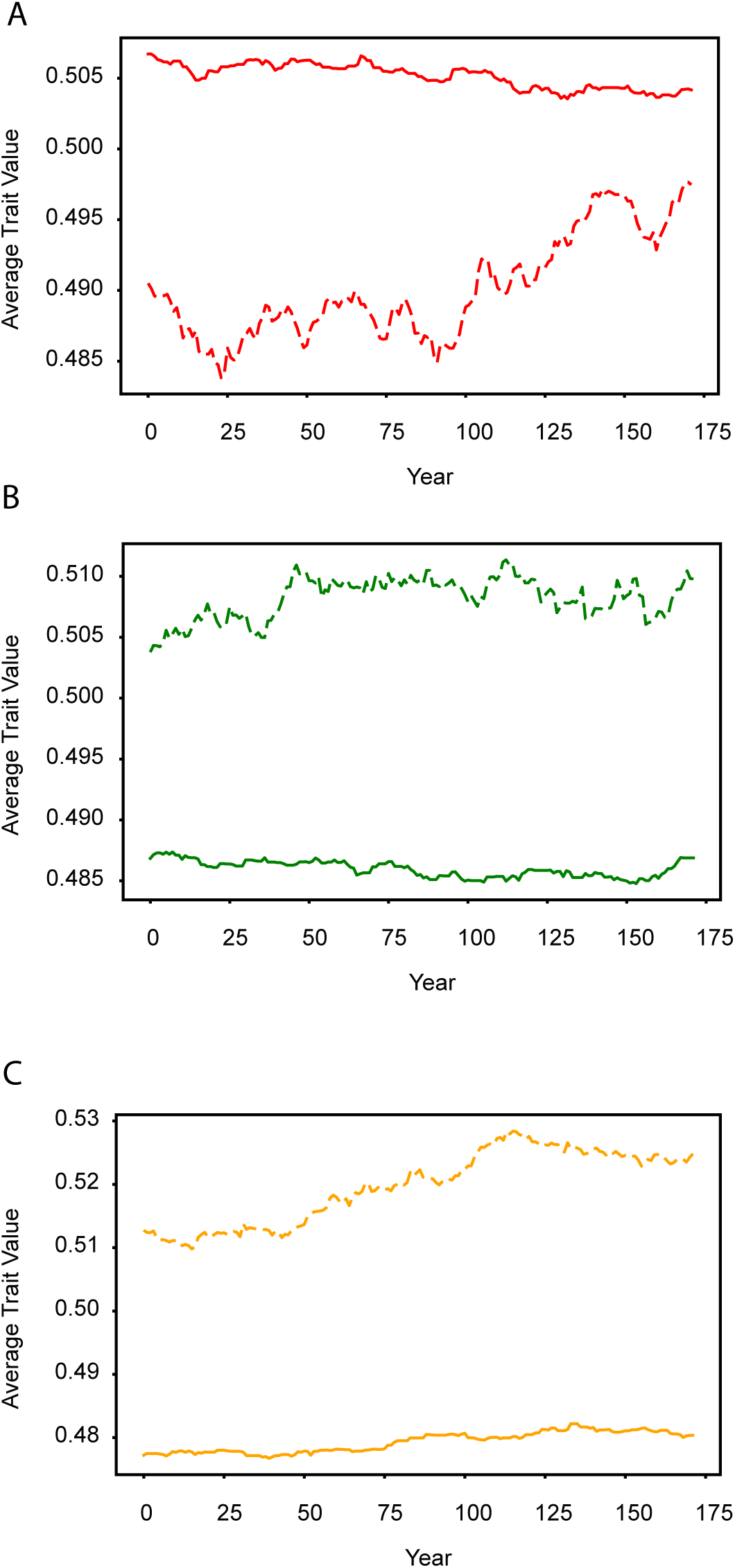
Impact of short, mid, and long interruptions on genetic and plasticity traits. (A) The evolution of genetic and plasticity traits for species experiencing short interruptions, where the interruption starts in year 10 and lasts for 25% of the total years. The plot highlights the impact of this short interruption on the trajectory of both traits. (B) The evolution of genetic and plasticity traits during mid interruptions, starting in year 20 and lasting for 50% of the total years. The figure shows how the traits are affected throughout the mid-length interruption period. (C) The evolution of genetic and plasticity traits during long interruptions, beginning in year 30 and continuing for 75% of the total years. The plot illustrates the extended impact of the long interruption on the traits over time.

### (c) Insights from sensitivity analysis

The sensitivity analysis of fitness across different cultivation scenarios, as modeled by the TEFI Model, reveals that the adaptation rate is the most influential factor under continuous cultivation, with a firstorder sensitivity index of 0.750 and a total-order sensitivity of 1.004 (Fig. 5A, 5B). This strong influence indicates that slow but steady adaptation primarily drives fitness in the absence of interruptions. Other parameters, such as mutation rate (first-order: 0.028, total-order: 0.261) and plasticity rate (first-order: 0.034, total-order: 0.265), contribute to fitness through interactions, but their direct effects are smaller. In the short interruption scenario, the plasticity effect becomes dominant (first-order: 0.058, total-order: 0.933), highlighting the role of plasticity in enabling rapid adaptation during brief interruptions (Fig. 5A, 5B). Recovery rate also plays a significant role (first-order: 0.040, total-order: 1.041), which aligns with the observed quick recovery of fitness post-interruption. In mid interruptions, the sensitivity of plasticity rate (first-order: 0.026, total-order: 0.930) and plasticity effect (first-order: 0.025, total-order: 0.950) reflects their importance in adapting to environmental stress, although the complex interactions between parameters lead to a fitness collapse after the interruption. For long interruptions, no parameter shows a strong direct influence on fitness, with the plasticity effect having the largest first-order sensitivity (0.037), but the cumulative effects of multiple parameters (total-order sensitivities all above 0.94) explain the severe fitness reduction and poor recovery in this scenario (Fig. 5A, 5B). In the noncontinuous interruption scenario, plasticity effect (first-order: 0.054, total-order: 1.026) drives fitness recovery, indicating that intermittent recovery periods during environmental disturbances allow populations to maintain relatively higher fitness levels.

**Figure 5:**
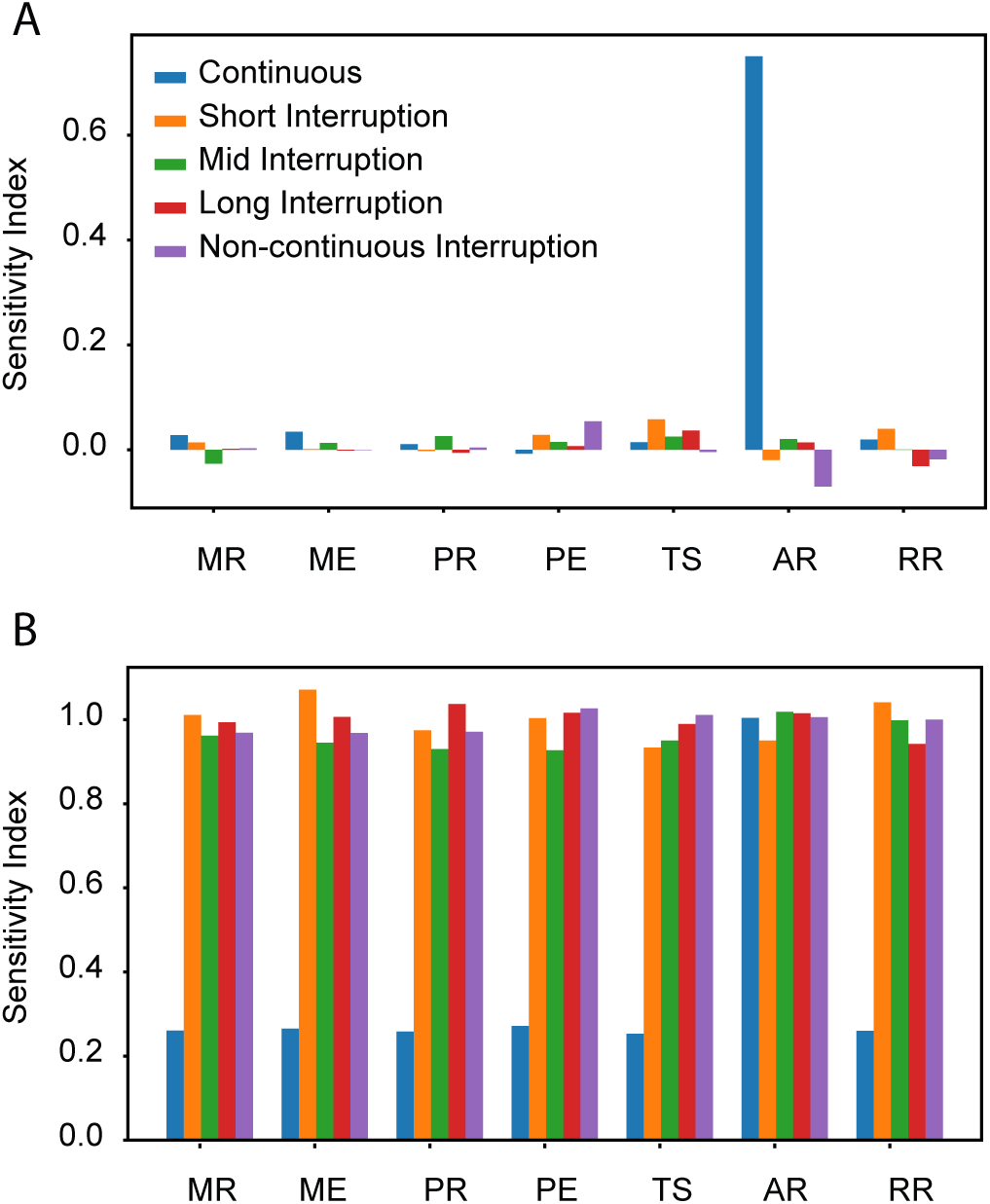
Sensitivity indices for fitness across cultivation and interruption scenarios. The first-order and total-order sensitivity indices (S1 and ST) for fitness in different cultivation scenarios. The bar plots in (A) illustrate the first-order sensitivity indices, showing the contribution of each parameter to fitness variation across continuous, short, mid, long, and non-continuous interruption scenarios. The parameters analyzed include mutation rate (MR), mutation effect (ME), plasticity rate (PR), plasticity effect (PE), temperature sensitivity (TS), adaptation rate (AR), and recovery rate (RR). In (B), the total-order sensitivity indices are plotted, revealing the overall impact of each parameter on fitness variation in the same cultivation scenarios.

The sensitivity analysis for average plasticity trait values also highlights the importance of plasticity effect and recovery rate in shaping trait evolution across different scenarios. Under continuous cultivation, the recovery rate has the highest first-order sensitivity (0.061) and total-order sensitivity (0.972), suggesting that populations in stable environments rely on recovery mechanisms to maintain plastic traits (Fig. 6A, 6B). In short interruptions, the plasticity effect (first-order: 0.041, total-order: 0.963) dominates, reflecting the need for rapid adaptation during and after brief disruptions. During mid interruptions, the plasticity effect (first-order: 0.086, total-order: 1.048) again plays a significant role, along with the plasticity rate (first-order: 0.043), as organisms adapt to extended periods of stress (Fig. 6A, 6B). For long interruptions, the plasticity effect (first-order: 0.064, total-order: 1.066) remains the most sensitive parameter, indicating that plastic traits undergo substantial changes under prolonged environmental pressures, even though fitness recovery is limited. In non-continuous interruptions, the plasticity effect (first-order: 0.090, total-order: 1.047) once again emerges as the key factor influencing plasticity traits, but the slight negative trends observed suggest that irregular disruptions may eventually limit the population’s adaptive flexibility.

**Figure 6:**
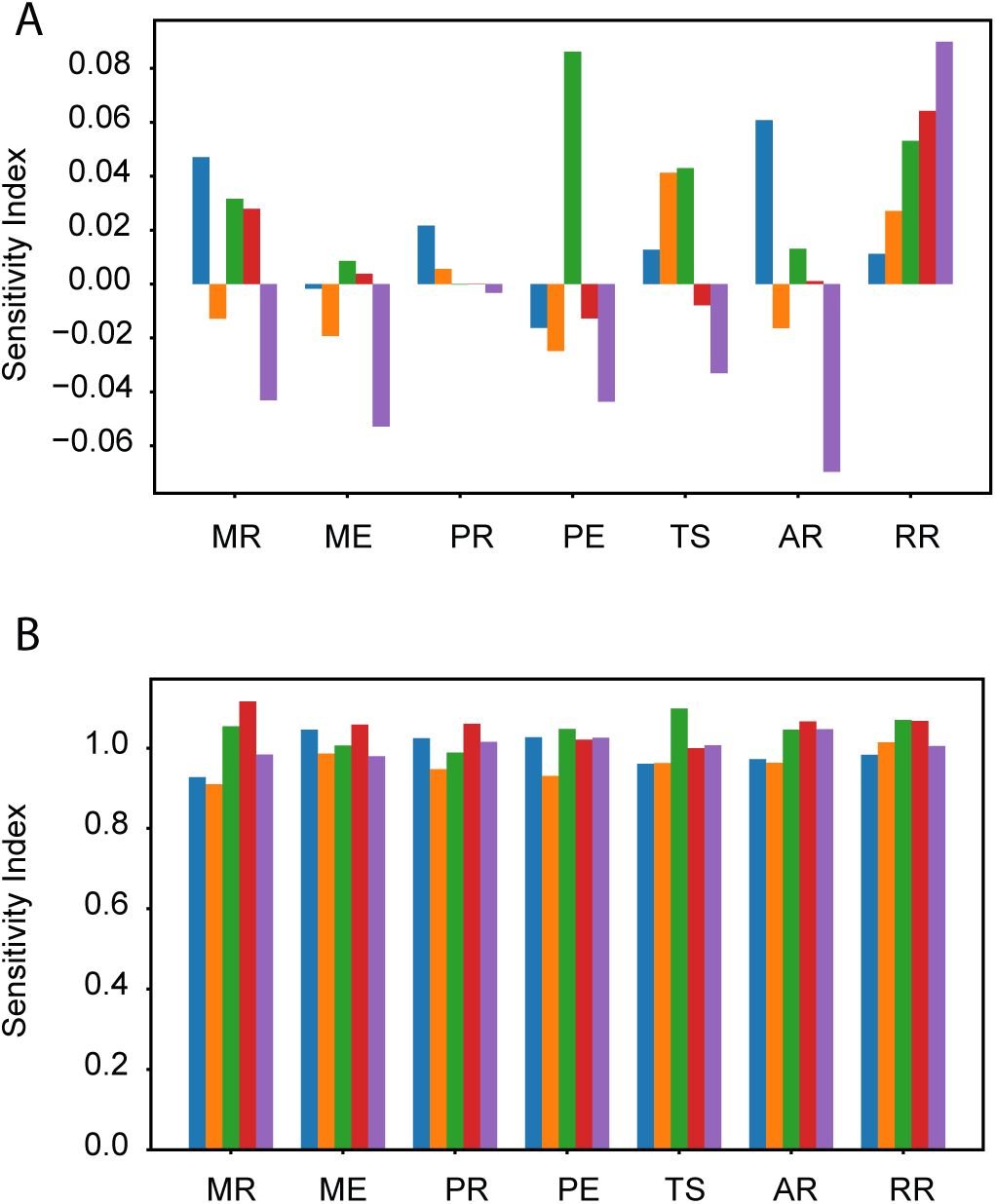
Sensitivity indices for plasticity traits across cultivation and interruption scenarios. The first-order and total-order sensitivity indices (S1 and ST) for plasticity traits across the same cultivation scenarios. The bar plots in (A) display the first-order sensitivity indices, highlighting the contribution of each parameter to plasticity trait variation in continuous, short, mid, long, and non-continuous interruption scenarios. Similarly, (B) presents the total-order sensitivity indices for plasticity traits, showing the overall impact of each parameter on plasticity trait variation in these scenarios.

### (d) Insights from case-studies

In rice and wheat, phenotypic plasticity was generally lower across traits, aligning with the theoretical continuous cultivation scenario, where populations experience minimal environmental disruption, leading to a gradual reduction in plastic traits. For rice, the overall median plasticity score was low (median = 0.04), with the highest plasticity observed in leaf blade width (median = 0.1116; IQR = 0.1035), while traits such as days to heading had much lower plasticity (median = 0.0108; IQR = 0.0122) (Fig. 7A). Similarly, wheat exhibited slightly higher overall plasticity (median = 0.11), with grain yield showing the highest plasticity (median = 0.2833; IQR = 0.0865), while plant height exhibited limited variability (median = 0.1039; IQR = 0.0223) (Fig. 7B). This reduction in plasticity aligns with theoretical predictions that under continuous cultivation, populations undergo minimal or gradual reductions in plastic traits over time. The stable environment allows for slow adaptation, where reduced plasticity ensures more consistent agricultural fitness, with fewer drastic changes required to maintain productivity. Theoretical insights suggest that in such scenarios, plasticity decreases as populations become well-adapted to stable conditions, minimizing the need for frequent phenotypic adjustments.

**Figure 7:**
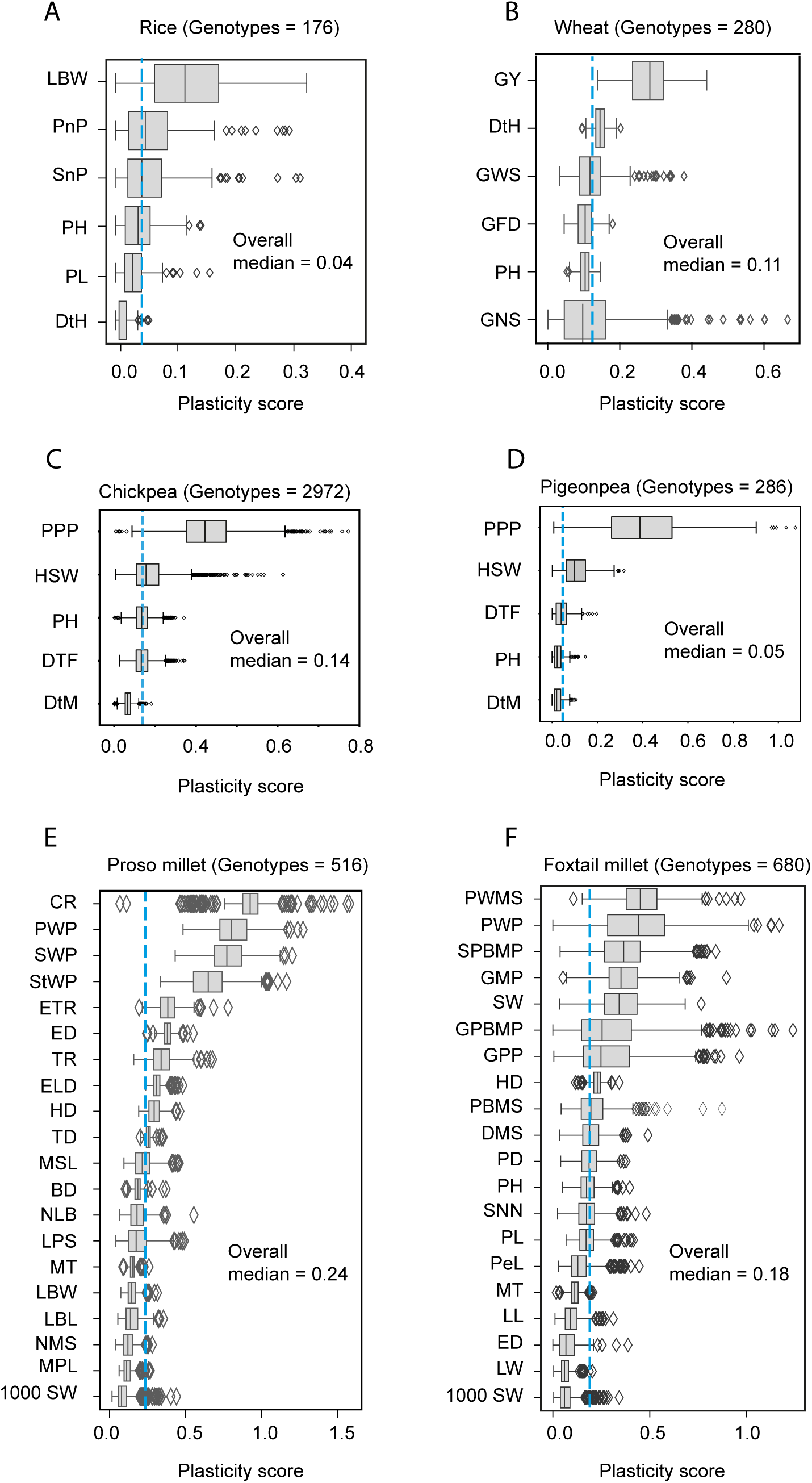
Comparative plasticity of traits in multiple crop species. Boxplots represent the plasticity scores of various traits measured across genotypes of four crop species: (A) Rice, (B) Wheat, (C) Chickpea, (D) Pigeonpea, (D) Proso millet, and (E) Foxtail millet. The overall median plasticity score is provided for each species (vertical blue line).

In line with the theoretical framework of continuous versus interrupted cultivation, chickpea shows moderate plasticity across traits (overall median = 0.14), suggesting adaptability under conditions with some environmental variability (Fig. 7C). This aligns with scenarios where moderate interruptions in cultivation drive the need for flexible responses. In chickpea, pods per plant demonstrates the highest plasticity (median = 0.444; IQR = 0.194), indicating an ability to adjust reproductive output, which can be advantageous in optimizing yield under variable conditions. Traits like days to flowering and plant height also show moderate plasticity (DTF: median = 0.132, IQR = 0.057; PH: median = 0.134, IQR = 0.052), while days to maturity exhibits lower plasticity (median = 0.066, IQR = 0.026), pointing to a balance between adaptability and stable maturation timing. Hundred-seed weight displays moderate plasticity (median = 0.156, IQR = 0.108), allowing flexibility in seed size as a response to environmental shifts. Pigeonpea exhibits lower overall plasticity (overall median = 0.05), aligning more closely with theoretical predictions for continuous cultivation scenarios, where stability is favored (Fig. 7D). Pods per plant stands out with relatively higher plasticity (median = 0.388; IQR = 0.267), suggesting adaptability in reproductive traits, possibly as a strategy for yield maintenance. Other traits, such as days to flowering and days to maturity, exhibit lower plasticity (DTF: median = 0.040, IQR = 0.046; DTM: median = 0.020, IQR = 0.028), as does plant height (PH) (median = 0.022, IQR = 0.027), reflecting a tendency toward stability. Hundred-seed weight shows moderate plasticity (median = 0.104; IQR = 0.088), offering some flexibility in seed size without extensive trait adjustment.

In contrast, broomcorn millet and foxtail millet displayed significantly higher plasticity, consistent with the theoretical interrupted cultivation scenarios, where populations face environmental stress and disruptions. In broomcorn millet, plasticity was particularly high across several traits (overall median = 0.24), especially in chaff rate (median = 0.9256; IQR = 0.1000) and panicle weight per plant (median = 0.8034; IQR = 0.1771), indicating strong adaptive responses to environmental variability ((Fig. 7E)). Foxtail millet exhibited moderate plasticity (overall median = 0.18), with traits like panicle weight per plant (median = 0.4410; IQR = 0.2929) and grain weight of the main panicle (median = 0.3514; IQR = 0.1477) being highly plastic (Fig. 7F). However, high plasticity has limits, as noted both in the theory and here in the results. While millets demonstrate adaptability to environmental stress, this elevated plasticity could lead to a reduction in agricultural fitness under prolonged or severe interruptions. The theoretical insights predict that although plasticity enables populations to survive environmental disruptions, extended reliance on high plasticity—especially in interrupted scenarios—can lead to fitness collapse if populations cannot recover adequately post-interruption. This could result in millets facing trade-offs in reproductive success and yield stability, as seen in traits like chaff rate and panicle weight per plant. Thus, while high plasticity is initially beneficial in dealing with environmental changes, both theory and this empirical data indicate that too much plasticity can limit the long-term recovery and stability of agricultural fitness.

## Discussions

Traditional approaches to modeling fitness in plant breeding and ecology, such as Genotype × Environment (G×E) interaction models, reaction norm models, process-based crop growth models, and extended rreeder’s equation for trait evolution, offer insights into crop adaptation, but lack certain features that the TEFI model incorporates. For example, G×E Interaction Models often represent fitness as *F* = *G* + *E* + (*G* × *E*), capturing direct genotype-environment interactions but without accounting for temporal adaptation. These models often utilize statistical tools like the additive main effects and multiplicative interaction (AMMI) model and GGE biplot analysis to dissect and visualize interactions. However, they primarily analyze performance data without explicitly modeling the underlying biological processes influencing fitness (Priyadarshan, 2019). Reaction norm models, such as the one used by Jarquín et al. (2014), describe phenotype as a function of both genetic and environmental variables, often through multiplicative terms that capture plastic responses to environmental changes. However, they generally omit any cumulative adaptation term over time, focusing instead on immediate genotype-by-environment interactions. By contrast, the TEFI model introduces an expanded fitness equation (Eq. 1), which integrates both immediate genotype-environment interactions and distinguishes between genetic and plastic trait contributions through separate summation terms 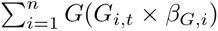 and 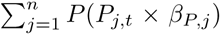. The model’s temporal adaptation term, *α* × *t* × *T_t_*, addsan evolutionary component, allowing it to capture gradual adaptation over time, unlike the more static frameworks of other models. In comparison, process-based models like GECROS integrate physiological processes with genotype-environment interactions to simulate short-term yield responses. However, they are typically limited to immediate yield predictions and do not fully capture the long-term evolution of fitness or adaptive genetic changes over time (Kadam et al., 2019; Resende et al., 2021). Similarly, the extended breeder’s equation has been adapted to consider environmental influences, enhancing the classical form, Δ*Z* = *h*^2^ × *S*, with additional terms that account for genotype-by-environment (G×E) interactions and the alignment between multi-environment trials (MET) and the target population of environments (TPE). This extension includes a genetic correlation term, *r_a_*(MET, TPE), to quantify the impact of MET-TPE alignment on trait evolution, thereby improving prediction accuracy under diverse environmental conditions (Cooper et al., 2023). However, despite these adaptations, the extended breeder’s equation does not incorporate the distinct separation of plastic and non-plastic trait contributions that the TEFI model offers, which enables a more detailed treatment of adaptive responses to environmental variation. In synthesizing elements from each of these approaches—environmental sensitivity, trait-specific contributions, and temporal adaptation—the TEFI model provides a novel, comprehensive tool to simulate crop fitness dynamics and long-term resilience under fluctuating environmental conditions. Phenotypic plasticity plays a key role in helping populations cope with environmental changes by enabling rapid phenotypic adjustments, which can delay extinction and allow time for genetic adaptation (Ashander et al., 2016; Reed et al., 2010). This temporary buffer, known as evolutionary rescue, allows populations to survive under new conditions while genetic evolution gradually aligns them with the new environmental norms (Lande, 2016; Fierst, 2011; Chevin et al., 2013). The findings from our study, as modeled through the TEFI Model, support this concept, as populations experiencing short interruptions showed rapid fitness recovery due to plastic traits, emphasizing plasticity’s role in quick adaptation to transient disturbances. Specifically, fitness increased during short interruptions (mean = 0.550; SD = 0.156), with a strong post-interruption recovery (mean = 0.886; SD = 0.189), illustrating how plasticity facilitates rapid adaptation when environmental stress is brief, as quantified by the TEFI Model. This contrasted with the slower, steady fitness progression seen in continuous cultivation (mean fitness = 0.497; SD = 0.149), where the minimal fitness change (slope = 0.000203) suggests gradual adaptation over time without major disruptions (Fig. 1 and 2). Therefore, short interruptions allow for faster fitness recovery due to the exploitation of plastic traits, while continuous cultivation fosters slower, genetically driven adaptation (Shaw and Etterson, 2012).

However, our results also show that the effectiveness of plasticity depends on environmental predictability, as seen in Scheiner et al. (2019) and Lande (2019). In highly unpredictable or prolonged interruptions—such as the mid and long interruption scenarios where fitness collapsed—plastic responses may become maladaptive, increasing the risk of extinction, as observed through simulations with the TEFI Model (Reed et al., 2010; Chevin et al., 2010). Specifically, under mid interruptions, fitness dropped from a relatively high pre-interruption level (mean = 0.492; SD = 0.150) to near total collapse post-interruption (mean = 0.0002; SD = 0.0015), and for long interruptions, fitness declined drastically during the interruption (mean = 0.083; SD = 0.087) and recovered only minimally afterward (mean = 0.012; SD = 0.033) (Fig. 1 and 2). These patterns suggest that cumulative impacts from extended environmental stress severely hinder population recovery, with fitness losses driven by a combination of reduced base fitness and diminished contributions from both genetic and plastic traits.

Lee et al. (2024) emphasize that plasticity’s role is often transient, with genetic accommodation taking over when environments stabilize, a pattern reflected in our results under continuous cultivation, where plasticity declined and genetic traits showed slow but steady change. In continuous cultivation, genetic traits exhibited a minimal positive rate of change (9.39 × 10*^−^*^6^), while plastic traits declined slightly (−7.06 × 10*^−^*^5^), suggesting that as the environment remained stable, populations relied more on genetic adaptation rather than plasticity (Fig. 3 and 4). This contrasts with non-continuous interruption scenarios, where intermittent environmental disturbances kept both plasticity and genetic traits active. Here, we observed a slight increase in genetic traits (1.52 × 10*^−^*^5^) while plastic traits declined (−6.71 × 10*^−^*^5^), suggesting that periodic interruptions allow genetic traits to adapt while plasticity decreases as populations stabilize between interruptions (Fig. 3 and 4). These findings underscore the dynamic interplay between plasticity and genetic evolution, where continuous cultivation favors slow genetic change and declining plasticity, while interrupted environments maintain the relevance of both plastic and genetic traits.

Additionally, maintaining plasticity comes with fitness costs, which can limit population persistence under rapidly changing conditions, as seen with high plasticity rates during extended interruptions (Chevin et al., 2010). Our sensitivity analysis revealed that in mid-length interruptions, plasticity rate and effect had significant total-order sensitivities (both ¿ 0.93), driving fitness dynamics under extended environmental stress (Fig. 5). The plasticity effect similarly dominated in short interruptions (first-order: 0.058, total-order: 0.933), allowing for rapid recovery after brief disruptions. Conversely, in long interruptions, the cumulative interaction of parameters (total-order sensitivities ¿ 0.94) drove the severe fitness reduction and poor recovery, highlighting that prolonged reliance on plasticity leads to diminishing returns and greater fitness loss. This aligns with theoretical expectations that, while plasticity is crucial for short-term adaptation, its fitness costs accumulate under extended or highly unpredictable conditions (De Lisle and Rowe, 2023; Bogan et al., 2024).

Our findings align with Lande (2019), who suggests that while plasticity is favored in fluctuating environments, its evolution is tempered by costs in stable conditions, where genetic traits offer better long-term fitness. In continuous cultivation, the adaptation rate had the highest first-order sensitivity (0.750) and total-order sensitivity (1.004), underscoring the importance of slow genetic adaptation in stable environments (Fig. 6). Meanwhile, in interrupted environments, plasticity effect and recovery rate drove fitness, especially in short interruptions where recovery rate (first-order: 0.040, total-order: 1.041) played a crucial role. This highlights a trade-off between plasticity and genetic traits, where plasticity dominates in unstable environments but genetic traits provide long-term stability in continuous cultivation.

Beyond ecological and evolutionary studies, similar principles of phenotypic plasticity apply in agricultural contexts, as seen in the Genomes to Fields (G2F) Maize project (Gage et al., 2017). The study highlights that modern breeding may have unintentionally reduced phenotypic plasticity in crops, particularly in intensively selected cultivars. Genomic regions selected during maize breeding were found to explain less G × E variation for yield, suggesting that the focus on improving stability in specific environments might have reduced the crop’s ability to adapt across diverse conditions. In our case studies of rice, wheat, broomcorn millet, and foxtail millet, similar plasticity trends were observed. Rice and wheat, both selected for stable environments, exhibited low plasticity (e.g., rice median plasticity = 0.04, wheat = 0.11), consistent with theoretical predictions of continuous cultivation (Fig. 7). In contrast, broomcorn millet and foxtail millet, which experience more environmental variability, showed higher plasticity (e.g., broomcorn millet median plasticity = 0.24, foxtail millet = 0.18), consistent with interrupted cultivation scenarios (Nimmo et al., 2023). Similarly, chickpea displayed moderate plasticity (overall median = 0.14), particularly in reproductive traits, indicating adaptability to certain environmental changes. Pigeonpea, however, exhibited lower plasticity (overall median = 0.05), reinforcing a tendency towards trait stability, possibly due to consistent growing conditions in its cultivated environments (Fig. 7). These findings reveal how different breeding targets—stability vs. adaptability—manifest in the plasticity and fitness trade-offs of various crops, with high plasticity aiding adaptation in unstable environments but potentially limiting long-term stability.

Overall, it is evident that phenotypic plasticity is often indirectly selected during breeding, even when breeders focus primarily on mean trait values. This suggests that intentionally selecting for high or low plasticity, alongside desirable mean phenotypes, could be challenging, especially when the genetic regulation of these traits is intertwined (Li et al., 2019). Moving forward, the key challenge will be finding ways to overcome these unintended trade-offs, particularly in efforts to breed for both environmental adaptability and high productivity.

Future research should focus on further investigating the genetic basis of plasticity and how it interacts with other fitness traits across different environmental conditions. Long-term studies on interrupted vs. continuous cultivation systems would help clarify the fitness trade-offs between plastic and genetic traits. Additionally, practical applications in crop breeding could explore how to balance plasticity with productivity, ensuring that crops maintain adaptability while also achieving high yields. A more detailed understanding of these dynamics will be essential for developing resilient crops in the face of climate variability.

## Supporting information

Additional Sheet 1, File S1, File S2, File S3

## Acknowledgements

This work received support from the Department of Biotechnology, India, under Grant No. BT/PR23613/BAP/118/354/2017, awarded to E.R. A.G. is the recipient of an IISER Tirupati Institutional PhD Fellowship.

## Author contributions

Ganesh Alagarasan: Conceptualization; data curation; formal analysis; investigation; methodology; software; validation; visualization; writing – original draft; writing – review and editing. Rajeev K. Varshney: chickpea and pigeonpea project; writing – review and editing. Eswarayya Ramireddy: Conceptualization; funding acquisition; methodology; supervision; writing – review and editing.

---

**Figure S1:**
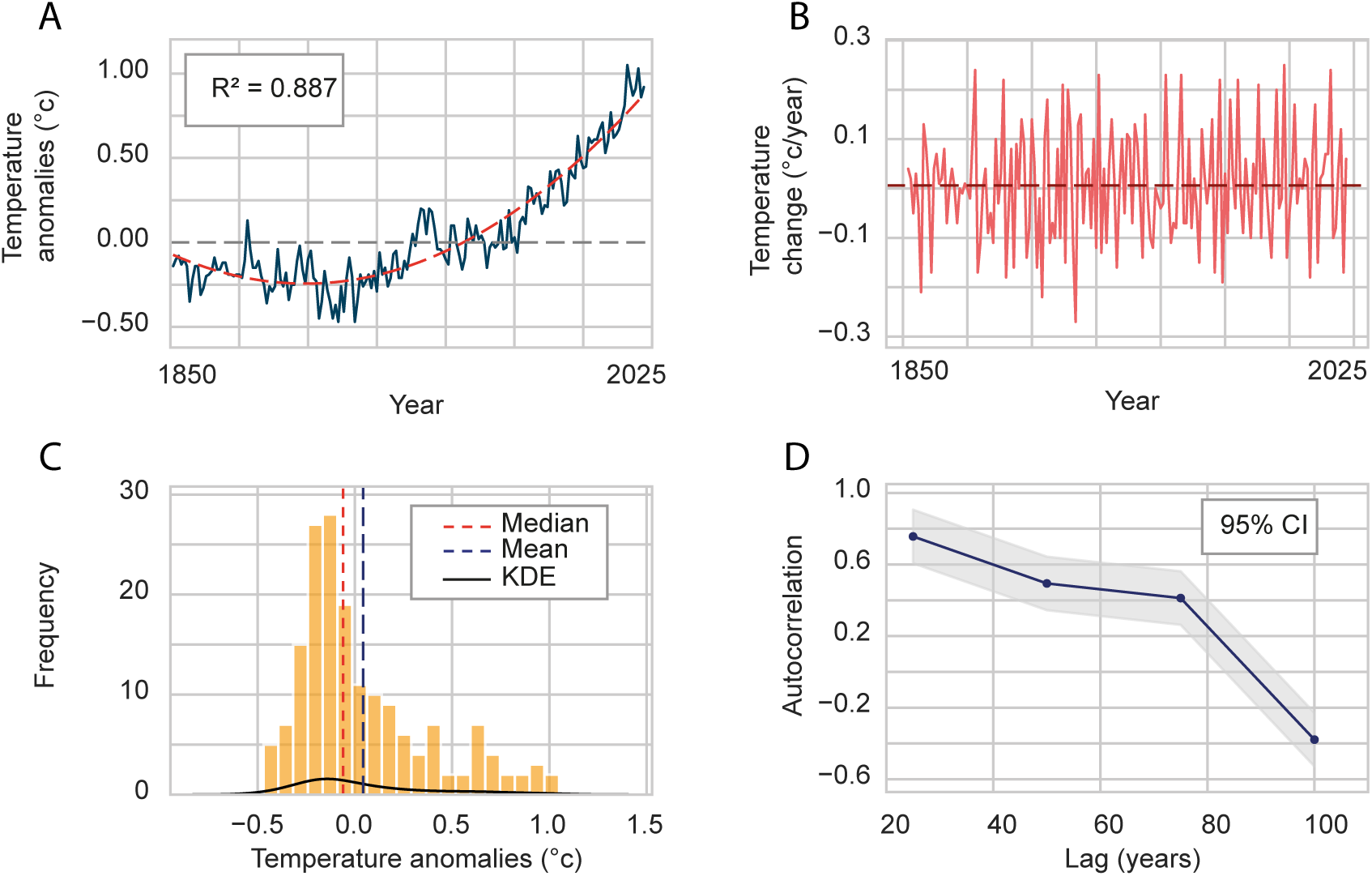
Characteristics of temperature anamolies. (A) Global surface mean annual temperature anomalies (°C) data, offering insights into climate trends through time, (B) annual rate of temperature change (°C year-1), (C) distribution and kernel density estimate (KDE) of mean annual temperature anomalies (°C), and (D) the autocorrelation of temperature data with different lag years.

