## Additional Sheet 1, File S1, File S2, File S3 for "Crops under continuous cultivation exhibit less plasticity than those under interrupted cultivation": File S1.py

```
import numpy as np
import pandas as pd
import matplotlib.pyplot as plt
import matplotlib
from scipy.stats import norm, linregress, ttest_ind, f_oneway

matplotlib.rcParams['svg.fonttype'] = 'none'
plt.rcParams.update({
    'font.size': 12, 
    'font.family': 'Arial',
    'font.weight': 'bold'
})

# Set seed for reproducibility
np.random.seed(42)

# Load temperature data
data = pd.read_csv('Temperature.csv')

# Define parameters
n_years = len(data)
temperature = data['Temperature'].values
n_regular = 50
n_intermittent = 50
n_traits = 5  # Number of genetic traits
n_plasticity_traits = 5  # Number of plasticity traits
mutation_rate = 0.005  # Probability of mutation per generation
mutation_effect = 0.05  # Effect of mutation on genetic traits
plasticity_rate = 0.01  # Rate at which plasticity traits change
plasticity_effect = 0.1  # Effect of plasticity on fitness
base_fitness = 0.5
temp_sensitivity = -0.02
adaptation_rate = 0.001
recovery_rate = 0.01  # Base rate at which fitness recovers after interruption

# Initialize genetic traits and plasticity traits
genetic_traits_regular = np.random.rand(n_regular, n_traits)
genetic_traits_intermittent_1 = np.random.rand(n_intermittent, n_traits)
genetic_traits_intermittent_2 = np.random.rand(n_intermittent, n_traits)
genetic_traits_intermittent_3 = np.random.rand(n_intermittent, n_traits)
genetic_traits_intermittent_non_continuous = np.random.rand(n_intermittent, n_traits)
plasticity_traits_regular = np.random.rand(n_regular, n_plasticity_traits)
plasticity_traits_intermittent_1 = np.random.rand(n_intermittent, n_plasticity_traits)
plasticity_traits_intermittent_2 = np.random.rand(n_intermittent, n_plasticity_traits)
plasticity_traits_intermittent_3 = np.random.rand(n_intermittent, n_plasticity_traits)
plasticity_traits_intermittent_non_continuous = np.random.rand(n_intermittent, n_plasticity_traits)

# Initialize a variable to accumulate the impact of interruption
cumulative_interruption_impact_1 = np.zeros(n_intermittent)
cumulative_interruption_impact_2 = np.zeros(n_intermittent)
cumulative_interruption_impact_3 = np.zeros(n_intermittent)
cumulative_interruption_impact_non_continuous = np.zeros(n_intermittent)

# Function to mutate only (for Non-Plastic / Genetic Traits)
def mutate_only(traits, mutation_rate, mutation_effect):
    mutation_mask = np.random.rand(*traits.shape) < mutation_rate
    mutations = np.random.normal(0, mutation_effect, traits.shape)
    new_traits = traits + mutation_mask * mutations
    return np.clip(new_traits, 0, 1)  # Ensuring trait values are between 0 and 1

# Function to mutate and plasticize (for Plastic / Plasticity Traits)
def mutate_and_plasticize(traits, mutation_rate, mutation_effect, plasticity_rate, plasticity_effect):
    mutation_mask = np.random.rand(*traits.shape) < mutation_rate
    mutations = np.random.normal(0, mutation_effect, traits.shape)
    plasticity_mask = np.random.rand(*traits.shape) < plasticity_rate
    plasticity_changes = np.random.normal(0, plasticity_effect, traits.shape)
    new_traits = traits + mutation_mask * mutations + plasticity_mask * plasticity_changes
    return np.clip(new_traits, 0, 1)  # Ensuring trait values are between 0 and 1

# Function to calculate fitness based on traits, temperature, and plasticity
def calculate_fitness(base_fitness, temp_sensitivity, adaptation_rate, temperature, year, genetic_traits, plasticity_traits, is_intermittent, interruption_params=None, cumulative_impact=None):
    fitness = base_fitness + temp_sensitivity * temperature + np.dot(genetic_traits, np.random.normal(0, 0.1, genetic_traits.shape[1]))
    fitness += np.dot(plasticity_traits, np.random.normal(0, 0.1, plasticity_traits.shape[1]))
    if not is_intermittent:
        fitness += adaptation_rate * year * temperature
    else:
        for start_year, end_year, reduction_factor, interruption_duration in interruption_params:
            if start_year <= year <= end_year:
                fitness -= reduction_factor  # Reduction during interruption
                cumulative_impact += reduction_factor * (end_year - start_year + 1)
            elif year > end_year:
                adjusted_base_fitness = base_fitness * (1 - cumulative_impact / 100)
                trait_influence = np.dot(genetic_traits, np.random.normal(0, 0.1, genetic_traits.shape[1])) * (1 - cumulative_impact / 100)
                plasticity_influence = np.dot(plasticity_traits, np.random.normal(0, 0.1, plasticity_traits.shape[1])) * (1 - cumulative_impact / 100)
                fitness = adjusted_base_fitness + temp_sensitivity * temperature + trait_influence + plasticity_influence
                adjusted_recovery_rate = recovery_rate / (1 + interruption_duration / 100) * (1 - fitness)
                recovery = adjusted_recovery_rate * (1 - fitness)  # Recovery after interruption
                fitness += recovery
    return np.clip(fitness, 0, 1)  # Ensuring fitness is between 0 and 1

# Random Sampling to Determine Non-Continuous Interruption Periods
n_interruption_periods = 3  # Number of random interruption periods
max_duration = 15  # Maximum duration for each interruption

non_continuous_periods = []
for _ in range(n_interruption_periods):
    start_year = np.random.randint(0, n_years - max_duration)
    end_year = start_year + np.random.randint(5, max_duration)  # Random duration between 5 and max_duration years
    reduction_factor = 0.3 * (temperature[end_year] - temperature[start_year])
    interruption_duration = end_year - start_year
    non_continuous_periods.append((start_year, end_year, reduction_factor, interruption_duration))

# Define interruption parameters for three types of interruptions: 25%, 50%, 75% of total years
short_duration = n_years // 4  # Short interruption: 25% of total years
mid_duration = n_years // 2    # Mid interruption: 50% of total years
long_duration = (3 * n_years) // 4  # Long interruption: 75% of total years

# Set start and end years for each type of interruption
short_start_year = 10
mid_start_year = 20
long_start_year = 30

short_end_year = short_start_year + short_duration
mid_end_year = mid_start_year + mid_duration
long_end_year = long_start_year + long_duration

# Calculate reduction factors based on temperature differences
short_reduction_factor = 0.5 * (temperature[short_end_year] - temperature[short_start_year])
mid_reduction_factor = 0.5 * (temperature[mid_end_year] - temperature[mid_start_year])
long_reduction_factor = 0.5 * (temperature[long_end_year] - temperature[long_start_year])

# Define interruption parameters for three types of interruptions
interruption_params_short = [(short_start_year, short_end_year, short_reduction_factor, short_duration)]
interruption_params_mid = [(mid_start_year, mid_end_year, mid_reduction_factor, mid_duration)]
interruption_params_long = [(long_start_year, long_end_year, long_reduction_factor, long_duration)]

# Fitness Before, During, and After Interruption (Intermittent Cultivation)
fitness_all_year = np.zeros((n_regular, n_years))
fitness_intermittent_short = np.zeros((n_intermittent, n_years))
fitness_intermittent_mid = np.zeros((n_intermittent, n_years))
fitness_intermittent_long = np.zeros((n_intermittent, n_years))
fitness_intermittent_non_continuous = np.zeros((n_intermittent, n_years))

# Simulate over the generations (years)
for year in range(n_years):
    # Update Non-Plastic / Genetic Traits with mutation only
    # These traits are genetic and do not undergo plastic changes.
    genetic_traits_regular = mutate_only(genetic_traits_regular, mutation_rate, mutation_effect)
    genetic_traits_intermittent_1 = mutate_only(genetic_traits_intermittent_1, mutation_rate, mutation_effect)
    genetic_traits_intermittent_2 = mutate_only(genetic_traits_intermittent_2, mutation_rate, mutation_effect)
    genetic_traits_intermittent_3 = mutate_only(genetic_traits_intermittent_3, mutation_rate, mutation_effect)
    genetic_traits_intermittent_non_continuous = mutate_only(genetic_traits_intermittent_non_continuous, mutation_rate, mutation_effect)

    # Update Plastic / Plasticity Traits with both mutation and plasticity changes
    # These traits are plastic and can adapt in response to the environment.
    plasticity_traits_regular = mutate_and_plasticize(plasticity_traits_regular, mutation_rate, mutation_effect, plasticity_rate, plasticity_effect)
    plasticity_traits_intermittent_1 = mutate_and_plasticize(plasticity_traits_intermittent_1, mutation_rate, mutation_effect, plasticity_rate, plasticity_effect)
    plasticity_traits_intermittent_2 = mutate_and_plasticize(plasticity_traits_intermittent_2, mutation_rate, mutation_effect, plasticity_rate, plasticity_effect)
    plasticity_traits_intermittent_3 = mutate_and_plasticize(plasticity_traits_intermittent_3, mutation_rate, mutation_effect, plasticity_rate, plasticity_effect)
    plasticity_traits_intermittent_non_continuous = mutate_and_plasticize(plasticity_traits_intermittent_non_continuous, mutation_rate, mutation_effect, plasticity_rate, plasticity_effect)

    # Calculate fitness for continuously-cultivated species
    fitness_all_year[:, year] = calculate_fitness(
        base_fitness, temp_sensitivity, adaptation_rate, temperature[year], year,
        genetic_traits_regular, plasticity_traits_regular, False
    )
    
    # Calculate fitness for intermittently-cultivated species (Short Interruption)
    fitness_intermittent_short[:, year] = calculate_fitness(
        base_fitness, temp_sensitivity, adaptation_rate, temperature[year], year,
        genetic_traits_intermittent_1, plasticity_traits_intermittent_1, True,
        interruption_params=interruption_params_short,
        cumulative_impact=cumulative_interruption_impact_1
    )
    
    # Calculate fitness for intermittently-cultivated species (Mid Interruption)
    fitness_intermittent_mid[:, year] = calculate_fitness(
        base_fitness, temp_sensitivity, adaptation_rate, temperature[year], year,
        genetic_traits_intermittent_2, plasticity_traits_intermittent_2, True,
        interruption_params=interruption_params_mid,
        cumulative_impact=cumulative_interruption_impact_2
    )
    
    # Calculate fitness for intermittently-cultivated species (Long Interruption)
    fitness_intermittent_long[:, year] = calculate_fitness(
        base_fitness, temp_sensitivity, adaptation_rate, temperature[year], year,
        genetic_traits_intermittent_3, plasticity_traits_intermittent_3, True,
        interruption_params=interruption_params_long,
        cumulative_impact=cumulative_interruption_impact_3
    )

    # Calculate fitness for intermittently-cultivated species (Non-continuous Interruption)
    fitness_intermittent_non_continuous[:, year] = calculate_fitness(
        base_fitness, temp_sensitivity, adaptation_rate, temperature[year], year,
        genetic_traits_intermittent_non_continuous, plasticity_traits_intermittent_non_continuous, True,
        interruption_params=non_continuous_periods,
        cumulative_impact=cumulative_interruption_impact_non_continuous
    )

# Average Rate of Change
avg_fitness_continuous = np.mean(fitness_all_year, axis=0)
avg_fitness_intermittent_short = np.mean(fitness_intermittent_short, axis=0)
avg_fitness_intermittent_mid = np.mean(fitness_intermittent_mid, axis=0)
avg_fitness_intermittent_long = np.mean(fitness_intermittent_long, axis=0)
avg_fitness_intermittent_non_continuous = np.mean(fitness_intermittent_non_continuous, axis=0)

# 1. Fitness Trajectory Without Interruption
plt.figure(figsize=(10, 6))
plt.plot(range(n_years), avg_fitness_continuous, label='Continuous Cultivation (No Interruption)', color='blue')
plt.xlabel('Year')
plt.ylabel('Average Fitness')
plt.title('Fitness Trajectory of Continuous Cultivation Without Interruption')
plt.legend()
plt.savefig('E:/New phytologist theory paper/Fig1A.svg', format='svg')
plt.show()

# 2. Fitness Before, During, and After Interruption (Non-continuous Interruption)
plt.figure(figsize=(10, 6))
for (start, end, _, _) in non_continuous_periods:
    plt.plot(range(n_years), avg_fitness_intermittent_non_continuous, label=f'Non-continuous Interruption {start}-{end}', linestyle='--', color='purple')
plt.xlabel('Year')
plt.ylabel('Average Fitness')
plt.title('Fitness During and After Non-continuous Interruption')
plt.legend()
plt.savefig('E:/New phytologist theory paper/Fig1B.svg', format='svg')
plt.show()

# 3. Fitness Before Interruption for Short, Mid, and Long Interruption
plt.figure(figsize=(10, 6))
plt.plot(range(short_start_year), avg_fitness_intermittent_short[:short_start_year], label='Intermittent Cultivation (Before Short Interruption)', color='red')
plt.plot(range(mid_start_year), avg_fitness_intermittent_mid[:mid_start_year], label='Intermittent Cultivation (Before Mid Interruption)', color='green')
plt.plot(range(long_start_year), avg_fitness_intermittent_long[:long_start_year], label='Intermittent Cultivation (Before Long Interruption)', color='orange')
plt.xlabel('Year')
plt.ylabel('Average Fitness')
plt.title('Fitness Before Interruption for Short, Mid, and Long Interruption')
plt.legend()
plt.savefig('E:/New phytologist theory paper/Fig1C.svg', format='svg')
plt.show()

# 4. Fitness During Interruption for All Types
plt.figure(figsize=(10, 6))
plt.plot(range(short_start_year, short_end_year + 1), avg_fitness_intermittent_short[short_start_year:short_end_year + 1], label='Intermittent Cultivation (During Short Interruption)', color='red')
plt.plot(range(mid_start_year, mid_end_year + 1), avg_fitness_intermittent_mid[mid_start_year:mid_end_year + 1], label='Intermittent Cultivation (During Mid Interruption)', color='green')
plt.plot(range(long_start_year, long_end_year + 1), avg_fitness_intermittent_long[long_start_year:long_end_year + 1], label='Intermittent Cultivation (During Long Interruption)', color='orange')
plt.xlabel('Year')
plt.ylabel('Average Fitness')
plt.title('Fitness During Interruption for Short, Mid, and Long Interruption')
plt.legend()
plt.savefig('E:/New phytologist theory paper/Fig1D.svg', format='svg')
plt.show()

# 5. Recovery Trajectory After Interruption for All Types
plt.figure(figsize=(10, 6))
plt.plot(range(short_end_year + 1, n_years), avg_fitness_intermittent_short[short_end_year + 1:], label='Intermittent Cultivation (Recovery After Short Interruption)', color='red')
plt.plot(range(mid_end_year + 1, n_years), avg_fitness_intermittent_mid[mid_end_year + 1:], label='Intermittent Cultivation (Recovery After Mid Interruption)', color='green')
plt.plot(range(long_end_year + 1, n_years), avg_fitness_intermittent_long[long_end_year + 1:], label='Intermittent Cultivation (Recovery After Long Interruption)', color='orange')
plt.xlabel('Year')
plt.ylabel('Average Fitness')
plt.title('Recovery Trajectory After Short, Mid, and Long Interruption')
plt.legend()
plt.savefig('E:/New phytologist theory paper/Fig1E.svg', format='svg')
plt.show()

# Statistical Analysis and Saving Results
stats_results = {
    'Statistic': [],
    'Value': []
}

# Calculate overall mean and standard deviation for each fitness trajectory
stats_results['Statistic'].extend(['Mean Fitness (Continuous)', 'Std Dev Fitness (Continuous)'])
stats_results['Value'].extend([
    np.mean(avg_fitness_continuous),
    np.std(avg_fitness_continuous)
])

# Add min and max values for continuous fitness
stats_results['Statistic'].extend(['Min Fitness (Continuous)', 'Max Fitness (Continuous)'])
stats_results['Value'].extend([
    np.min(avg_fitness_continuous),
    np.max(avg_fitness_continuous)
])

# Calculate slope of the fitness trajectory for continuous cultivation
slope, intercept, r_value, p_value, std_err = linregress(range(n_years), avg_fitness_continuous)
stats_results['Statistic'].append('Slope of Fitness Trajectory (Continuous)')
stats_results['Value'].append(slope)

# Calculate mean, std, min, max, and slope for intermittent interruptions (short, mid, long, non-continuous)
interruption_types = [
    ('Short Interruption', avg_fitness_intermittent_short),
    ('Mid Interruption', avg_fitness_intermittent_mid),
    ('Long Interruption', avg_fitness_intermittent_long),
    ('Non-continuous Interruption', avg_fitness_intermittent_non_continuous)
]

for label, fitness_data in interruption_types:
    # Before Interruption Stats
    stats_results['Statistic'].extend([
        f'Mean Fitness Before {label}',
        f'Std Dev Fitness Before {label}',
        f'Min Fitness Before {label}',
        f'Max Fitness Before {label}'
    ])
    stats_results['Value'].extend([
        np.mean(fitness_data[:short_start_year]),
        np.std(fitness_data[:short_start_year]),
        np.min(fitness_data[:short_start_year]),
        np.max(fitness_data[:short_start_year])
    ])

    # During Interruption Stats
    if label == 'Short Interruption':
        start_year, end_year = short_start_year, short_end_year
    elif label == 'Mid Interruption':
        start_year, end_year = mid_start_year, mid_end_year
    elif label == 'Long Interruption':
        start_year, end_year = long_start_year, long_end_year
    else:
        # Non-continuous interruption
        start_year, end_year = non_continuous_periods[0][0], non_continuous_periods[-1][1]
    
    stats_results['Statistic'].extend([
        f'Mean Fitness During {label}',
        f'Std Dev Fitness During {label}',
        f'Min Fitness During {label}',
        f'Max Fitness During {label}'
    ])
    stats_results['Value'].extend([
        np.mean(fitness_data[start_year:end_year + 1]),
        np.std(fitness_data[start_year:end_year + 1]),
        np.min(fitness_data[start_year:end_year + 1]),
        np.max(fitness_data[start_year:end_year + 1])
    ])

    # After Interruption Stats
    stats_results['Statistic'].extend([
        f'Mean Fitness After {label}',
        f'Std Dev Fitness After {label}',
        f'Min Fitness After {label}',
        f'Max Fitness After {label}'
    ])
    stats_results['Value'].extend([
        np.mean(fitness_data[end_year + 1:]),
        np.std(fitness_data[end_year + 1:]),
        np.min(fitness_data[end_year + 1:]),
        np.max(fitness_data[end_year + 1:])
    ])

# Statistical comparisons using ANOVA and T-tests
anova_result = f_oneway(avg_fitness_intermittent_short, avg_fitness_intermittent_mid, avg_fitness_intermittent_long, avg_fitness_intermittent_non_continuous)
stats_results['Statistic'].append('ANOVA p-value (Between Interruption Types)')
stats_results['Value'].append(anova_result.pvalue)

# T-tests between different interruption types
for i in range(len(interruption_types)):
    for j in range(i + 1, len(interruption_types)):
        label1, fitness_data1 = interruption_types[i]
        label2, fitness_data2 = interruption_types[j]
        t_stat, p_val = ttest_ind(fitness_data1, fitness_data2)
        stats_results['Statistic'].append(f'T-test p-value ({label1} vs {label2})')
        stats_results['Value'].append(p_val)

# Creating DataFrame to store results
stats_df = pd.DataFrame(stats_results)

# Save results to Excel file
stats_df.to_excel('E:/fitness_statistics_results.xlsx', index=False)

print("Statistical analysis completed and saved to 'fitness_statistics_results.xlsx'")
```
