## Additional Sheet 1, File S1, File S2, File S3 for "Crops under continuous cultivation exhibit less plasticity than those under interrupted cultivation": File S2.py

```
import numpy as np
import pandas as pd
import matplotlib.pyplot as plt
import matplotlib
from scipy.stats import norm, linregress, ttest_ind, f_oneway

matplotlib.rcParams['svg.fonttype'] = 'none'
plt.rcParams.update({
    'font.size': 12, 
    'font.family': 'Arial',
    'font.weight': 'bold'
})

# Set seed for reproducibility
np.random.seed(42)

# Load temperature data
data = pd.read_csv('Temperature.csv')

# Define parameters
n_years = len(data)
temperature = data['Temperature'].values
n_regular = 50
n_intermittent = 50
n_traits = 5  # Number of genetic traits
n_plasticity_traits = 5  # Number of plasticity traits
mutation_rate = 0.005  # Probability of mutation per generation
mutation_effect = 0.05  # Effect of mutation on genetic traits
plasticity_rate = 0.01  # Rate at which plasticity traits change
plasticity_effect = 0.1  # Effect of plasticity on fitness
base_fitness = 0.5
temp_sensitivity = -0.02
adaptation_rate = 0.001
recovery_rate = 0.01  # Base rate at which fitness recovers after interruption

# Initialize genetic traits and plasticity traits
genetic_traits_regular = np.random.rand(n_regular, n_traits)
genetic_traits_intermittent_1 = np.random.rand(n_intermittent, n_traits)
genetic_traits_intermittent_2 = np.random.rand(n_intermittent, n_traits)
genetic_traits_intermittent_3 = np.random.rand(n_intermittent, n_traits)
genetic_traits_intermittent_non_continuous = np.random.rand(n_intermittent, n_traits)
plasticity_traits_regular = np.random.rand(n_regular, n_plasticity_traits)
plasticity_traits_intermittent_1 = np.random.rand(n_intermittent, n_plasticity_traits)
plasticity_traits_intermittent_2 = np.random.rand(n_intermittent, n_plasticity_traits)
plasticity_traits_intermittent_3 = np.random.rand(n_intermittent, n_plasticity_traits)
plasticity_traits_intermittent_non_continuous = np.random.rand(n_intermittent, n_plasticity_traits)

# Initialize a variable to accumulate the impact of interruption
cumulative_interruption_impact_1 = np.zeros(n_intermittent)
cumulative_interruption_impact_2 = np.zeros(n_intermittent)
cumulative_interruption_impact_3 = np.zeros(n_intermittent)
cumulative_interruption_impact_non_continuous = np.zeros(n_intermittent)

# Function to mutate only (for Non-Plastic / Genetic Traits)
def mutate_only(traits, mutation_rate, mutation_effect):
    mutation_mask = np.random.rand(*traits.shape) < mutation_rate
    mutations = np.random.normal(0, mutation_effect, traits.shape)
    new_traits = traits + mutation_mask * mutations
    return np.clip(new_traits, 0, 1)  # Ensuring trait values are between 0 and 1

# Function to mutate and plasticize (for Plastic / Plasticity Traits)
def mutate_and_plasticize(traits, mutation_rate, mutation_effect, plasticity_rate, plasticity_effect):
    mutation_mask = np.random.rand(*traits.shape) < mutation_rate
    mutations = np.random.normal(0, mutation_effect, traits.shape)
    plasticity_mask = np.random.rand(*traits.shape) < plasticity_rate
    plasticity_changes = np.random.normal(0, plasticity_effect, traits.shape)
    new_traits = traits + mutation_mask * mutations + plasticity_mask * plasticity_changes
    return np.clip(new_traits, 0, 1)  # Ensuring trait values are between 0 and 1

# Function to calculate fitness based on traits, temperature, and plasticity
def calculate_fitness(base_fitness, temp_sensitivity, adaptation_rate, temperature, year, genetic_traits, plasticity_traits, is_intermittent, interruption_params=None, cumulative_impact=None):
    fitness = base_fitness + temp_sensitivity * temperature + np.dot(genetic_traits, np.random.normal(0, 0.1, genetic_traits.shape[1]))
    fitness += np.dot(plasticity_traits, np.random.normal(0, 0.1, plasticity_traits.shape[1]))
    if not is_intermittent:
        fitness += adaptation_rate * year * temperature
    else:
        for start_year, end_year, reduction_factor, interruption_duration in interruption_params:
            if start_year <= year <= end_year:
                fitness -= reduction_factor  # Reduction during interruption
                cumulative_impact += reduction_factor * (end_year - start_year + 1)
            elif year > end_year:
                adjusted_base_fitness = base_fitness * (1 - cumulative_impact / 100)
                trait_influence = np.dot(genetic_traits, np.random.normal(0, 0.1, genetic_traits.shape[1])) * (1 - cumulative_impact / 100)
                plasticity_influence = np.dot(plasticity_traits, np.random.normal(0, 0.1, plasticity_traits.shape[1])) * (1 - cumulative_impact / 100)
                fitness = adjusted_base_fitness + temp_sensitivity * temperature + trait_influence + plasticity_influence
                adjusted_recovery_rate = recovery_rate / (1 + interruption_duration / 100) * (1 - fitness)
                recovery = adjusted_recovery_rate * (1 - fitness)  # Recovery after interruption
                fitness += recovery
    return np.clip(fitness, 0, 1)  # Ensuring fitness is between 0 and 1

# Random Sampling to Determine Non-Continuous Interruption Periods
n_interruption_periods = 3  # Number of random interruption periods
max_duration = 15  # Maximum duration for each interruption

non_continuous_periods = []
for _ in range(n_interruption_periods):
    start_year = np.random.randint(0, n_years - max_duration)
    end_year = start_year + np.random.randint(5, max_duration)  # Random duration between 5 and max_duration years
    reduction_factor = 0.3 * (temperature[end_year] - temperature[start_year])
    interruption_duration = end_year - start_year
    non_continuous_periods.append((start_year, end_year, reduction_factor, interruption_duration))

# Define interruption parameters for three types of interruptions: 25%, 50%, 75% of total years
short_duration = n_years // 4  # Short interruption: 25% of total years
mid_duration = n_years // 2    # Mid interruption: 50% of total years
long_duration = (3 * n_years) // 4  # Long interruption: 75% of total years

# Set start and end years for each type of interruption
short_start_year = 10
mid_start_year = 20
long_start_year = 30

short_end_year = short_start_year + short_duration
mid_end_year = mid_start_year + mid_duration
long_end_year = long_start_year + long_duration

# Calculate reduction factors based on temperature differences
short_reduction_factor = 0.5 * (temperature[short_end_year] - temperature[short_start_year])
mid_reduction_factor = 0.5 * (temperature[mid_end_year] - temperature[mid_start_year])
long_reduction_factor = 0.5 * (temperature[long_end_year] - temperature[long_start_year])

# Define interruption parameters for three types of interruptions
interruption_params_short = [(short_start_year, short_end_year, short_reduction_factor, short_duration)]
interruption_params_mid = [(mid_start_year, mid_end_year, mid_reduction_factor, mid_duration)]
interruption_params_long = [(long_start_year, long_end_year, long_reduction_factor, long_duration)]

# Fitness Before, During, and After Interruption (Intermittent Cultivation)
fitness_all_year = np.zeros((n_regular, n_years))
fitness_intermittent_short = np.zeros((n_intermittent, n_years))
fitness_intermittent_mid = np.zeros((n_intermittent, n_years))
fitness_intermittent_long = np.zeros((n_intermittent, n_years))
fitness_intermittent_non_continuous = np.zeros((n_intermittent, n_years))

# Arrays to track average genetic and plasticity trait values over time
avg_genetic_regular = np.zeros((n_years, n_traits))
avg_plastic_regular = np.zeros((n_years, n_plasticity_traits))
avg_genetic_intermittent_short = np.zeros((n_years, n_traits))
avg_plastic_intermittent_short = np.zeros((n_years, n_plasticity_traits))
avg_genetic_intermittent_mid = np.zeros((n_years, n_traits))
avg_plastic_intermittent_mid = np.zeros((n_years, n_plasticity_traits))
avg_genetic_intermittent_long = np.zeros((n_years, n_traits))
avg_plastic_intermittent_long = np.zeros((n_years, n_plasticity_traits))
avg_genetic_intermittent_non_continuous = np.zeros((n_years, n_traits))
avg_plastic_intermittent_non_continuous = np.zeros((n_years, n_plasticity_traits))

# Simulate over the generations (years)
for year in range(n_years):
    # Update Non-Plastic / Genetic Traits with mutation only
    genetic_traits_regular = mutate_only(genetic_traits_regular, mutation_rate, mutation_effect)
    genetic_traits_intermittent_1 = mutate_only(genetic_traits_intermittent_1, mutation_rate, mutation_effect)
    genetic_traits_intermittent_2 = mutate_only(genetic_traits_intermittent_2, mutation_rate, mutation_effect)
    genetic_traits_intermittent_3 = mutate_only(genetic_traits_intermittent_3, mutation_rate, mutation_effect)
    genetic_traits_intermittent_non_continuous = mutate_only(genetic_traits_intermittent_non_continuous, mutation_rate, mutation_effect)

    # Update Plastic / Plasticity Traits with both mutation and plasticity changes
    plasticity_traits_regular = mutate_and_plasticize(plasticity_traits_regular, mutation_rate, mutation_effect, plasticity_rate, plasticity_effect)
    plasticity_traits_intermittent_1 = mutate_and_plasticize(plasticity_traits_intermittent_1, mutation_rate, mutation_effect, plasticity_rate, plasticity_effect)
    plasticity_traits_intermittent_2 = mutate_and_plasticize(plasticity_traits_intermittent_2, mutation_rate, mutation_effect, plasticity_rate, plasticity_effect)
    plasticity_traits_intermittent_3 = mutate_and_plasticize(plasticity_traits_intermittent_3, mutation_rate, mutation_effect, plasticity_rate, plasticity_effect)
    plasticity_traits_intermittent_non_continuous = mutate_and_plasticize(plasticity_traits_intermittent_non_continuous, mutation_rate, mutation_effect, plasticity_rate, plasticity_effect)

    # Track average genetic and plastic traits for each group
    avg_genetic_regular[year] = np.mean(genetic_traits_regular, axis=0)
    avg_plastic_regular[year] = np.mean(plasticity_traits_regular, axis=0)
    avg_genetic_intermittent_short[year] = np.mean(genetic_traits_intermittent_1, axis=0)
    avg_plastic_intermittent_short[year] = np.mean(plasticity_traits_intermittent_1, axis=0)
    avg_genetic_intermittent_mid[year] = np.mean(genetic_traits_intermittent_2, axis=0)
    avg_plastic_intermittent_mid[year] = np.mean(plasticity_traits_intermittent_2, axis=0)
    avg_genetic_intermittent_long[year] = np.mean(genetic_traits_intermittent_3, axis=0)
    avg_plastic_intermittent_long[year] = np.mean(plasticity_traits_intermittent_3, axis=0)
    avg_genetic_intermittent_non_continuous[year] = np.mean(genetic_traits_intermittent_non_continuous, axis=0)
    avg_plastic_intermittent_non_continuous[year] = np.mean(plasticity_traits_intermittent_non_continuous, axis=0)

    # Calculate fitness for continuously-cultivated species
    fitness_all_year[:, year] = calculate_fitness(
        base_fitness, temp_sensitivity, adaptation_rate, temperature[year], year,
        genetic_traits_regular, plasticity_traits_regular, False
    )
    
    # Calculate fitness for intermittently-cultivated species (Short Interruption)
    fitness_intermittent_short[:, year] = calculate_fitness(
        base_fitness, temp_sensitivity, adaptation_rate, temperature[year], year,
        genetic_traits_intermittent_1, plasticity_traits_intermittent_1, True,
        interruption_params=interruption_params_short,
        cumulative_impact=cumulative_interruption_impact_1
    )

    # Calculate fitness for intermittently-cultivated species (Mid Interruption)
    fitness_intermittent_mid[:, year] = calculate_fitness(
        base_fitness, temp_sensitivity, adaptation_rate, temperature[year], year,
        genetic_traits_intermittent_2, plasticity_traits_intermittent_2, True,
        interruption_params=interruption_params_mid,
        cumulative_impact=cumulative_interruption_impact_2
    )

    # Calculate fitness for intermittently-cultivated species (Long Interruption)
    fitness_intermittent_long[:, year] = calculate_fitness(
        base_fitness, temp_sensitivity, adaptation_rate, temperature[year], year,
        genetic_traits_intermittent_3, plasticity_traits_intermittent_3, True,
        interruption_params=interruption_params_long,
        cumulative_impact=cumulative_interruption_impact_3
    )

    # Calculate fitness for intermittently-cultivated species (Non-continuous Interruption)
    fitness_intermittent_non_continuous[:, year] = calculate_fitness(
        base_fitness, temp_sensitivity, adaptation_rate, temperature[year], year,
        genetic_traits_intermittent_non_continuous, plasticity_traits_intermittent_non_continuous, True,
        interruption_params=non_continuous_periods,
        cumulative_impact=cumulative_interruption_impact_non_continuous
    )

# Plot evolution of genetic vs plasticity traits for continuous cultivation
plt.figure(figsize=(10, 6))
plt.plot(range(n_years), np.mean(avg_genetic_regular, axis=1), label='Average Genetic Traits (Continuous)', linestyle='-', color='blue')
plt.plot(range(n_years), np.mean(avg_plastic_regular, axis=1), label='Average Plasticity Traits (Continuous)', linestyle='--', color='blue')
plt.xlabel('Year')
plt.ylabel('Average Trait Value')
plt.title('Evolution of Genetic vs Plasticity Traits Over Time (Continuous Cultivation)')
plt.legend
plt.savefig('E:/New phytologist theory paper/Fig2A.svg', format='svg')
plt.show()

# Plot evolution of genetic vs plasticity traits for short interruption
plt.figure(figsize=(10, 6))
plt.plot(range(n_years), np.mean(avg_genetic_intermittent_short, axis=1), label='Average Genetic Traits (Short Interruption)', linestyle='-', color='red')
plt.plot(range(n_years), np.mean(avg_plastic_intermittent_short, axis=1), label='Average Plasticity Traits (Short Interruption)', linestyle='--', color='red')
plt.xlabel('Year')
plt.ylabel('Average Trait Value')
plt.title('Evolution of Genetic vs Plasticity Traits Over Time (Short Interruption)')
plt.legend()
plt.savefig('E:/New phytologist theory paper/Fig2B.svg', format='svg')
plt.show()

# Plot evolution of genetic vs plasticity traits for mid interruption
plt.figure(figsize=(10, 6))
plt.plot(range(n_years), np.mean(avg_genetic_intermittent_mid, axis=1), label='Average Genetic Traits (Mid Interruption)', linestyle='-', color='green')
plt.plot(range(n_years), np.mean(avg_plastic_intermittent_mid, axis=1), label='Average Plasticity Traits (Mid Interruption)', linestyle='--', color='green')
plt.xlabel('Year')
plt.ylabel('Average Trait Value')
plt.title('Evolution of Genetic vs Plasticity Traits Over Time (Mid Interruption)')
plt.legend()
plt.savefig('E:/New phytologist theory paper/Fig2C.svg', format='svg')
plt.show()

# Plot evolution of genetic vs plasticity traits for long interruption
plt.figure(figsize=(10, 6))
plt.plot(range(n_years), np.mean(avg_genetic_intermittent_long, axis=1), label='Average Genetic Traits (Long Interruption)', linestyle='-', color='orange')
plt.plot(range(n_years), np.mean(avg_plastic_intermittent_long, axis=1), label='Average Plasticity Traits (Long Interruption)', linestyle='--', color='orange')
plt.xlabel('Year')
plt.ylabel('Average Trait Value')
plt.title('Evolution of Genetic vs Plasticity Traits Over Time (Long Interruption)')
plt.legend()
plt.savefig('E:/New phytologist theory paper/Fig2D.svg', format='svg')
plt.show()

# Plot evolution of genetic vs plasticity traits for non-continuous interruption
plt.figure(figsize=(10, 6))
plt.plot(range(n_years), np.mean(avg_genetic_intermittent_non_continuous, axis=1), label='Average Genetic Traits (Non-continuous Interruption)', linestyle='-', color='purple')
plt.plot(range(n_years), np.mean(avg_plastic_intermittent_non_continuous, axis=1), label='Average Plasticity Traits (Non-continuous Interruption)', linestyle='--', color='purple')
plt.xlabel('Year')
plt.ylabel('Average Trait Value')
plt.title('Evolution of Genetic vs Plasticity Traits Over Time (Non-continuous Interruption)')
plt.legend()
plt.savefig('E:/New phytologist theory paper/Fig2E.svg', format='svg')
plt.show()

# Calculate rate of change for genetic vs plastic traits using linear regression
slope_genetic_regular, _, _, _, _ = linregress(range(n_years), np.mean(avg_genetic_regular, axis=1))
slope_plastic_regular, _, _, _, _ = linregress(range(n_years), np.mean(avg_plastic_regular, axis=1))
print(f"Rate of Change (Slope) for Genetic Traits (Continuous): {slope_genetic_regular}")
print(f"Rate of Change (Slope) for Plastic Traits (Continuous): {slope_plastic_regular}")

slope_genetic_short, _, _, _, _ = linregress(range(n_years), np.mean(avg_genetic_intermittent_short, axis=1))
slope_plastic_short, _, _, _, _ = linregress(range(n_years), np.mean(avg_plastic_intermittent_short, axis=1))
print(f"Rate of Change (Slope) for Genetic Traits (Short Interruption): {slope_genetic_short}")
print(f"Rate of Change (Slope) for Plastic Traits (Short Interruption): {slope_plastic_short}")

slope_genetic_mid, _, _, _, _ = linregress(range(n_years), np.mean(avg_genetic_intermittent_mid, axis=1))
slope_plastic_mid, _, _, _, _ = linregress(range(n_years), np.mean(avg_plastic_intermittent_mid, axis=1))
print(f"Rate of Change (Slope) for Genetic Traits (Mid Interruption): {slope_genetic_mid}")
print(f"Rate of Change (Slope) for Plastic Traits (Mid Interruption): {slope_plastic_mid}")

slope_genetic_long, _, _, _, _ = linregress(range(n_years), np.mean(avg_genetic_intermittent_long, axis=1))
slope_plastic_long, _, _, _, _ = linregress(range(n_years), np.mean(avg_plastic_intermittent_long, axis=1))
print(f"Rate of Change (Slope) for Genetic Traits (Long Interruption): {slope_genetic_long}")
print(f"Rate of Change (Slope) for Plastic Traits (Long Interruption): {slope_plastic_long}")

slope_genetic_non_continuous, _, _, _, _ = linregress(range(n_years), np.mean(avg_genetic_intermittent_non_continuous, axis=1))
slope_plastic_non_continuous, _, _, _, _ = linregress(range(n_years), np.mean(avg_plastic_intermittent_non_continuous, axis=1))
print(f"Rate of Change (Slope) for Genetic Traits (Non-continuous Interruption): {slope_genetic_non_continuous}")
print(f"Rate of Change (Slope) for Plastic Traits (Non-continuous Interruption): {slope_plastic_non_continuous}")
```
