## Additional Sheet 1, File S1, File S2, File S3 for "Crops under continuous cultivation exhibit less plasticity than those under interrupted cultivation": File S3.py

```
import numpy as np
import pandas as pd
import matplotlib.pyplot as plt
from scipy.stats import sem, t, ttest_ind, pearsonr, gaussian_kde
import matplotlib
from SALib.sample import saltelli
from SALib.analyze import sobol

matplotlib.rcParams['svg.fonttype'] = 'none'
plt.rcParams.update({
    'font.size': 12, 
    'font.family': 'Arial',
    'font.weight': 'bold'
})

# Set seed for reproducibility
np.random.seed(42)

# Load temperature data
data = pd.read_csv('Temperature.csv')

# Define parameters
n_years = len(data)
temperature = data['Temperature'].values
n_regular = 50
n_intermittent = 50
n_traits = 5  # Number of genetic traits
n_plasticity_traits = 5  # Number of plasticity traits
base_fitness = 0.5


# Define interruption parameters
short_duration = n_years // 4
mid_duration = n_years // 2
long_duration = (3 * n_years) // 4

short_start_year = 10
mid_start_year = 20
long_start_year = 30

short_end_year = short_start_year + short_duration
mid_end_year = mid_start_year + mid_duration
long_end_year = long_start_year + long_duration

short_reduction_factor = 0.5 * (temperature[short_end_year] - temperature[short_start_year])
mid_reduction_factor = 0.5 * (temperature[mid_end_year] - temperature[mid_start_year])
long_reduction_factor = 0.5 * (temperature[long_end_year] - temperature[long_start_year])

# Define interruption parameters
interruption_params_short = [(short_start_year, short_end_year, short_reduction_factor, short_duration)]
interruption_params_mid = [(mid_start_year, mid_end_year, mid_reduction_factor, mid_duration)]
interruption_params_long = [(long_start_year, long_end_year, long_reduction_factor, long_duration)]

# Generate non-continuous interruption periods
n_interruption_periods = 3  # Number of random interruption periods
max_duration = 15  # Maximum duration for each interruption

non_continuous_periods = []
for _ in range(n_interruption_periods):
    start_year = np.random.randint(0, n_years - max_duration)
    end_year = start_year + np.random.randint(5, max_duration)  # Random duration between 5 and max_duration years
    reduction_factor = 0.3 * (temperature[end_year] - temperature[start_year])
    interruption_duration = end_year - start_year
    non_continuous_periods.append((start_year, end_year, reduction_factor, interruption_duration))

# Function to simulate mutation (for genetic traits only)
def mutate(traits, mutation_rate, mutation_effect):
    mutation_mask = np.random.rand(*traits.shape) < mutation_rate
    mutations = np.random.normal(0, mutation_effect, traits.shape)
    new_traits = traits + mutation_mask * mutations
    return np.clip(new_traits, 0, 1)  # Ensuring trait values are between 0 and 1

# Function to simulate mutation and plasticity changes (for plasticity traits only)
def mutate_and_plasticize(traits, mutation_rate, mutation_effect, plasticity_rate, plasticity_effect):
    mutation_mask = np.random.rand(*traits.shape) < mutation_rate
    mutations = np.random.normal(0, mutation_effect, traits.shape)
    plasticity_mask = np.random.rand(*traits.shape) < plasticity_rate
    plasticity_changes = np.random.normal(0, plasticity_effect, traits.shape)
    new_traits = traits + mutation_mask * mutations + plasticity_mask * plasticity_changes
    return np.clip(new_traits, 0, 1)  # Ensuring trait values are between 0 and 1

# Function to calculate fitness based on traits, temperature, and plasticity
def calculate_fitness(base_fitness, temp_sensitivity, adaptation_rate, temperature, year, genetic_traits, plasticity_traits, is_intermittent, interruption_params=None, cumulative_impact=None):
    fitness = base_fitness + temp_sensitivity * temperature + np.dot(genetic_traits, np.random.normal(0, 0.1, genetic_traits.shape[1]))  # Genetic trait influence
    fitness += np.dot(plasticity_traits, np.random.normal(0, 0.1, plasticity_traits.shape[1]))  # Plasticity influence
    if not is_intermittent:
        fitness += adaptation_rate * year * temperature  # Continuous adaptation
    else:
        for start_year, end_year, reduction_factor, interruption_duration in interruption_params:
            if year < start_year:
                pass  # Same fitness as regular before interruption
            elif start_year <= year <= end_year:
                fitness -= reduction_factor  # Reduction during interruption
                cumulative_impact += reduction_factor * (end_year - start_year + 1)
            else:
                # Adjust base fitness and trait influence based on cumulative interruption impact
                adjusted_base_fitness = base_fitness * (1 - cumulative_impact / 100)
                trait_influence = np.dot(genetic_traits, np.random.normal(0, 0.1, genetic_traits.shape[1])) * (1 - cumulative_impact / 100)
                plasticity_influence = np.dot(plasticity_traits, np.random.normal(0, 0.1, plasticity_traits.shape[1])) * (1 - cumulative_impact / 100)
                fitness = adjusted_base_fitness + temp_sensitivity * temperature + trait_influence + plasticity_influence
                adjusted_recovery_rate = recovery_rate / (1 + interruption_duration / 100) * (1 - fitness)
                recovery = adjusted_recovery_rate * (1 - fitness)  # Recovery after interruption
                fitness += recovery
    return np.clip(fitness, 0, 1)  # Ensuring fitness is between 0 and 1

# Sobol Sensitivity Analysis Setup
problem = {
    'num_vars': 7,
    'names': ['mutation_rate', 'mutation_effect', 'plasticity_rate', 'plasticity_effect', 'temp_sensitivity', 'adaptation_rate', 'recovery_rate'],
    'bounds': [[0.001, 0.01], [0.01, 0.1], [0.001, 0.05], [0.05, 0.2], [-0.05, 0.01], [0.0001, 0.005], [0.001, 0.02]]
}

# Generate parameter samples using Saltelli's method
param_values = saltelli.sample(problem, 1000)

# Storage for model outputs for each scenario
Y_continuous = []
Y_short = []
Y_mid = []
Y_long = []
Y_non_continuous = []

# Storage for average plasticity trait values
P_continuous = []
P_short = []
P_mid = []
P_long = []
P_non_continuous = []

# Run the model for each parameter set
for i, params in enumerate(param_values):
    mutation_rate, mutation_effect, plasticity_rate, plasticity_effect, temp_sensitivity, adaptation_rate,recovery_rate = params
    
    # Initialize traits for each scenario
    genetic_traits_continuous = np.random.rand(n_regular, n_traits)
    genetic_traits_short = np.random.rand(n_intermittent, n_traits)
    genetic_traits_mid = np.random.rand(n_intermittent, n_traits)
    genetic_traits_long = np.random.rand(n_intermittent, n_traits)
    genetic_traits_non_continuous = np.random.rand(n_intermittent, n_traits)
    
    plasticity_traits_continuous = np.random.rand(n_regular, n_plasticity_traits)
    plasticity_traits_short = np.random.rand(n_intermittent, n_plasticity_traits)
    plasticity_traits_mid = np.random.rand(n_intermittent, n_plasticity_traits)
    plasticity_traits_long = np.random.rand(n_intermittent, n_plasticity_traits)
    plasticity_traits_non_continuous = np.random.rand(n_intermittent, n_plasticity_traits)
    
    # Initialize cumulative interruption impacts
    cumulative_impact_short = np.zeros(n_intermittent)
    cumulative_impact_mid = np.zeros(n_intermittent)
    cumulative_impact_long = np.zeros(n_intermittent)
    cumulative_impact_non_continuous = np.zeros(n_intermittent)
    
    # Define lists to store fitness and plasticity values for each year
    fitness_continuous = []
    fitness_short = []
    fitness_mid = []
    fitness_long = []
    fitness_non_continuous = []
    
    avg_plasticity_continuous = []
    avg_plasticity_short = []
    avg_plasticity_mid = []
    avg_plasticity_long = []
    avg_plasticity_non_continuous = []

    for year in range(n_years):
        # Continuous Cultivation
        genetic_traits_continuous = mutate(genetic_traits_continuous, mutation_rate, mutation_effect)
        plasticity_traits_continuous = mutate_and_plasticize(plasticity_traits_continuous, mutation_rate, mutation_effect, plasticity_rate, plasticity_effect)
        fitness_value_continuous = calculate_fitness(
            base_fitness, temp_sensitivity, adaptation_rate, temperature[year], year,
            genetic_traits_continuous, plasticity_traits_continuous, False
        )
        fitness_continuous.append(np.mean(fitness_value_continuous))
        avg_plasticity_continuous.append(np.mean(plasticity_traits_continuous))
        
        # Short Interruption
        genetic_traits_short = mutate(genetic_traits_short, mutation_rate, mutation_effect)
        plasticity_traits_short = mutate_and_plasticize(plasticity_traits_short, mutation_rate, mutation_effect, plasticity_rate, plasticity_effect)
        fitness_value_short = calculate_fitness(
            base_fitness, temp_sensitivity, adaptation_rate, temperature[year], year,
            genetic_traits_short, plasticity_traits_short, True,
            interruption_params=interruption_params_short,
            cumulative_impact=cumulative_impact_short
        )
        fitness_short.append(np.mean(fitness_value_short))
        avg_plasticity_short.append(np.mean(plasticity_traits_short))
        
        # Mid Interruption
        genetic_traits_mid = mutate(genetic_traits_mid, mutation_rate, mutation_effect)
        plasticity_traits_mid = mutate_and_plasticize(plasticity_traits_mid, mutation_rate, mutation_effect, plasticity_rate, plasticity_effect)
        fitness_value_mid = calculate_fitness(
            base_fitness, temp_sensitivity, adaptation_rate, temperature[year], year,
            genetic_traits_mid, plasticity_traits_mid, True,
            interruption_params=interruption_params_mid,
            cumulative_impact=cumulative_impact_mid
        )
        fitness_mid.append(np.mean(fitness_value_mid))
        avg_plasticity_mid.append(np.mean(plasticity_traits_mid))
        
        # Long Interruption
        genetic_traits_long = mutate(genetic_traits_long, mutation_rate, mutation_effect)
        plasticity_traits_long = mutate_and_plasticize(plasticity_traits_long, mutation_rate, mutation_effect, plasticity_rate, plasticity_effect)
        fitness_value_long = calculate_fitness(
            base_fitness, temp_sensitivity, adaptation_rate, temperature[year], year,
            genetic_traits_long, plasticity_traits_long, True,
            interruption_params=interruption_params_long,
            cumulative_impact=cumulative_impact_long
        )
        fitness_long.append(np.mean(fitness_value_long))
        avg_plasticity_long.append(np.mean(plasticity_traits_long))
        
        # Non-continuous Interruption
        genetic_traits_non_continuous = mutate(genetic_traits_non_continuous, mutation_rate, mutation_effect)
        plasticity_traits_non_continuous = mutate_and_plasticize(plasticity_traits_non_continuous, mutation_rate, mutation_effect, plasticity_rate, plasticity_effect)
        fitness_value_non_continuous = calculate_fitness(
            base_fitness, temp_sensitivity, adaptation_rate, temperature[year], year,
            genetic_traits_non_continuous, plasticity_traits_non_continuous, True,
            interruption_params=non_continuous_periods,
            cumulative_impact=cumulative_impact_non_continuous
        )
        fitness_non_continuous.append(np.mean(fitness_value_non_continuous))
        avg_plasticity_non_continuous.append(np.mean(plasticity_traits_non_continuous))
    
    # Calculate average fitness and plasticity across all years for each scenario and store for sensitivity analysis
    Y_continuous.append(np.mean(fitness_continuous))
    Y_short.append(np.mean(fitness_short))
    Y_mid.append(np.mean(fitness_mid))
    Y_long.append(np.mean(fitness_long))
    Y_non_continuous.append(np.mean(fitness_non_continuous))

    P_continuous.append(np.mean(avg_plasticity_continuous))
    P_short.append(np.mean(avg_plasticity_short))
    P_mid.append(np.mean(avg_plasticity_mid))
    P_long.append(np.mean(avg_plasticity_long))
    P_non_continuous.append(np.mean(avg_plasticity_non_continuous))

# Perform Sobol analysis for fitness
Si_fitness_continuous = sobol.analyze(problem, np.array(Y_continuous))
Si_fitness_short = sobol.analyze(problem, np.array(Y_short))
Si_fitness_mid = sobol.analyze(problem, np.array(Y_mid))
Si_fitness_long = sobol.analyze(problem, np.array(Y_long))
Si_fitness_non_continuous = sobol.analyze(problem, np.array(Y_non_continuous))

# Perform Sobol analysis for plasticity traits
Si_plasticity_continuous = sobol.analyze(problem, np.array(P_continuous))
Si_plasticity_short = sobol.analyze(problem, np.array(P_short))
Si_plasticity_mid = sobol.analyze(problem, np.array(P_mid))
Si_plasticity_long = sobol.analyze(problem, np.array(P_long))
Si_plasticity_non_continuous = sobol.analyze(problem, np.array(P_non_continuous))

# Print first-order and total-order sensitivity indices for fitness in each scenario
print("Fitness - Continuous Cultivation Scenario:")
print("First-order sensitivity indices:", Si_fitness_continuous['S1'])
print("Total-order sensitivity indices:", Si_fitness_continuous['ST'])

print("Fitness - Short Interruption Scenario:")
print("First-order sensitivity indices:", Si_fitness_short['S1'])
print("Total-order sensitivity indices:", Si_fitness_short['ST'])

print("Fitness - Mid Interruption Scenario:")
print("First-order sensitivity indices:", Si_fitness_mid['S1'])
print("Total-order sensitivity indices:", Si_fitness_mid['ST'])

print("Fitness - Long Interruption Scenario:")
print("First-order sensitivity indices:", Si_fitness_long['S1'])
print("Total-order sensitivity indices:", Si_fitness_long['ST'])

print("Fitness - Non-continuous Interruption Scenario:")
print("First-order sensitivity indices:", Si_fitness_non_continuous['S1'])
print("Total-order sensitivity indices:", Si_fitness_non_continuous['ST'])

# Print first-order and total-order sensitivity indices for plasticity traits in each scenario
print("Plasticity - Continuous Cultivation Scenario:")
print("First-order sensitivity indices:", Si_plasticity_continuous['S1'])
print("Total-order sensitivity indices:", Si_plasticity_continuous['ST'])

print("Plasticity - Short Interruption Scenario:")
print("First-order sensitivity indices:", Si_plasticity_short['S1'])
print("Total-order sensitivity indices:", Si_plasticity_short['ST'])

print("Plasticity - Mid Interruption Scenario:")
print("First-order sensitivity indices:", Si_plasticity_mid['S1'])
print("Total-order sensitivity indices:", Si_plasticity_mid['ST'])

print("Plasticity - Long Interruption Scenario:")
print("First-order sensitivity indices:", Si_plasticity_long['S1'])
print("Total-order sensitivity indices:", Si_plasticity_long['ST'])

print("Plasticity - Non-continuous Interruption Scenario:")
print("First-order sensitivity indices:", Si_plasticity_non_continuous['S1'])
print("Total-order sensitivity indices:", Si_plasticity_non_continuous['ST'])

# Extract Sobol indices for plotting (Fitness and Plasticity)
scenarios = ['Continuous', 'Short Interruption', 'Mid Interruption', 'Long Interruption', 'Non-continuous Interruption']
S1_fitness_indices = [Si_fitness_continuous['S1'], Si_fitness_short['S1'], Si_fitness_mid['S1'], Si_fitness_long['S1'], Si_fitness_non_continuous['S1']]
ST_fitness_indices = [Si_fitness_continuous['ST'], Si_fitness_short['ST'], Si_fitness_mid['ST'], Si_fitness_long['ST'], Si_fitness_non_continuous['ST']]

S1_plasticity_indices = [Si_plasticity_continuous['S1'], Si_plasticity_short['S1'], Si_plasticity_mid['S1'], Si_plasticity_long['S1'], Si_plasticity_non_continuous['S1']]
ST_plasticity_indices = [Si_plasticity_continuous['ST'], Si_plasticity_short['ST'], Si_plasticity_mid['ST'], Si_plasticity_long['ST'], Si_plasticity_non_continuous['ST']]

params = problem['names']

# Plotting first-order indices for fitness
fig, axs = plt.subplots(2, 1, figsize=(10, 12))

for i, scenario in enumerate(scenarios):
    axs[0].bar(np.arange(len(params)) + i * 0.15, S1_fitness_indices[i], width=0.15, label=scenario)

axs[0].set_title('First-order Sensitivity Indices (S1) - Fitness')
axs[0].set_xlabel('Parameters')
axs[0].set_ylabel('Sensitivity Index')
axs[0].set_xticks(np.arange(len(params)) + 0.3)
axs[0].set_xticklabels(params)
axs[0].legend()

# Plotting total-order indices for fitness
for i, scenario in enumerate(scenarios):
    axs[1].bar(np.arange(len(params)) + i * 0.15, ST_fitness_indices[i], width=0.15, label=scenario)

axs[1].set_title('Total-order Sensitivity Indices (ST) - Fitness')
axs[1].set_xlabel('Parameters')
axs[1].set_ylabel('Sensitivity Index')
axs[1].set_xticks(np.arange(len(params)) + 0.3)
axs[1].set_xticklabels(params)
axs[1].legend()

plt.tight_layout()
plt.savefig('E:/New phytologist theory paper/Fig3AB.svg', format='svg')
plt.show()

# Plotting first-order indices for plasticity traits
fig, axs = plt.subplots(2, 1, figsize=(10, 12))

for i, scenario in enumerate(scenarios):
    axs[0].bar(np.arange(len(params)) + i * 0.15, S1_plasticity_indices[i], width=0.15, label=scenario)

axs[0].set_title('First-order Sensitivity Indices (S1) - Plasticity Traits')
axs[0].set_xlabel('Parameters')
axs[0].set_ylabel('Sensitivity Index')
axs[0].set_xticks(np.arange(len(params)) + 0.3)
axs[0].set_xticklabels(params)
axs[0].legend()

# Plotting total-order indices for plasticity traits
for i, scenario in enumerate(scenarios):
    axs[1].bar(np.arange(len(params)) + i * 0.15, ST_plasticity_indices[i], width=0.15, label=scenario)

axs[1].set_title('Total-order Sensitivity Indices (ST) - Plasticity Traits')
axs[1].set_xlabel('Parameters')
axs[1].set_ylabel('Sensitivity Index')
axs[1].set_xticks(np.arange(len(params)) + 0.3)
axs[1].set_xticklabels(params)
axs[1].legend()

plt.tight_layout()
plt.savefig('E:/New phytologist theory paper/Fig3CD.svg', format='svg')
plt.show()
```
